# Pre-implantation non-steroidal anti-inflammatory drug treatment disrupts mouse embryo spacing and post-implantation chamber morphogenesis

**DOI:** 10.1101/2025.11.02.686054

**Authors:** Noura Massri, Savannah E. LaBuda, Michelle Oltean, Ripla Arora

## Abstract

Non-steroidal anti-inflammatory drugs (NSAIDs), which inhibit the prostaglandin synthase (PTGS) enzymes PTGS1 and PTGS2, may present a risk when consumed during early pregnancy. The impact of PTGS enzyme inhibition by NSAIDS on the three-dimensional organization and function of uterine compartments in context of implantation success is not well known. Here we show that pre-implantation treatment of mouse pregnancy with indomethacin, an NSAID that inhibits both PTGS1 and PTGS2, results in embryo crowding, delayed embryo implantation, defective implantation chamber formation that eventually results in significant pregnancy loss. These effects are dose dependent as lower doses of indomethacin do not show these effects on pregnancy. When embryo spacing defects were observed, a significant proportion of embryos that showed delayed growth or resorption were located in distinct decidual sites, suggesting that additional mechanisms beyond spatial distribution are at play. These findings highlight the potential risks associated with NSAID use during early pregnancy and underscore the need for further research into the underlying molecular mechanisms and the development of safer pain management strategies during pregnancy.

## INTRODUCTION

Non-steroidal anti-inflammatory drugs (NSAIDs), include drugs such as indomethacin, aspirin, naproxen, and ibuprofen and represent an important class of medications frequently prescribed for alleviating pain and inflammation (Funk, 2001). NSAIDs inhibit prostaglandin synthase enzyme (PTGS)1 and/or PTGS2 (Black et al., 2019) to reduce the levels of circulating prostaglandins (PGs) that are the cause of pain. Although NSAIDs provide effective pain relief for many pregnant women, population-based studies investigating NSAID use and adverse pregnancy outcomes have yielded conflicting results (Black et al., 2019). While some studies suggest that NSAID consumption during the early stages of pregnancy may be linked to an increased risk of miscarriage (Li et al., 2003; Nakhai-Pour et al., 2011; Nielsen et al., 2001) other studies have found no association (Daniel et al., 2014; Edwards et al., 2012). These inconsistent findings are complicated by methodological limitations, such as inadequate exposure measurement and a failure to consider over-the-counter NSAID use. Thus, understanding the risks of NSAID use during early pregnancy is essential for healthcare providers and expectant mothers to make informed pain management decisions during this critical fetal development period.

Prostaglandins are important lipid mediators involved in signaling during reproductive processes. In early pregnancy, they coordinate various functions, including ovulation, fertilization, embryo transport, implantation, immune cell recruitment, vascular remodeling, and decidualization. (Kennedy et al., 2007; Lim et al., 1999; Lim et al., 1997). At the maternal-fetal interface, specific PG subtypes play distinctive roles: PGE2 promotes uterine receptivity and implantation, PGF2α regulates uterine contractility, prostacyclin (PGI2) is critical for transformation of endometrial stromal cells into decidual cells that support implantation and early placental development (Kennedy and Zamecnik, 1978; Lim et al., 1999). In mice, PGI2 is the most abundant PG at implantation sites, over 4 times higher than PG, and mediates embryo implantation via the PPARδ receptor (Lim et al., 1999). Furthermore, receptors for PGs, DP and EP3, cooperate to promote decidualization (Sakamoto et al., 2024). The coordinated production of PGs facilitated by both PTGS1 and PTGS2 enzymes, creates the optimal uterine environment for successful embryo implantation, and overall pregnancy success (Aikawa et al., 2024; Chakraborty et al., 1996; Massri and Arora, 2025; Sugimoto et al., 2025).

Given the ethical considerations in studying human subjects, rodent models have become essential for addressing NSAID risks. Inhibition of PG networks by NSAIDs can significantly impact pregnancy establishment and maintenance in humans and rodents (Black et al., 2019; Kennedy et al., 2007; Reese et al., 2000; Reese et al., 2001). In rodents, NSAIDs affect multiple aspects of pregnancy, including ovulation, embryo implantation, decidualization, duration of gestation, and litter size (Aiken, 1972; Diao et al., 2007; Kennedy et al., 2007; Reese et al., 2000; Reese et al., 2001). In mice, early pregnancy proceeds through precisely timed stages: fertilization around gestational day (GD) 0.5, blastocyst development by GD3 (Miller, 2018; Wang and Dey, 2006), entry of blastocysts into uterus and attachment to the uterine epithelium from GD3 to GD4 (Flores et al., 2020), formation of a V-shaped implantation chamber by GD4.5 (Madhavan et al., 2022), stromal decidualization and extensive vascular remodeling through GD7.5 (Kim et al., 2013; Marchetto et al., 2020; Massri et al., 2023). NSAIDs demonstrate timing- and dose-dependent effects on implantation. The peri-implantation window (GD0-GD4) represents a vulnerable period; in mice, indomethacin (PTGS1 and PTGS2 inhibitor) treatment on GD0-3 caused dose and timing-dependent pregnancy termination (Lau et al., 1973). In rats indomethacin administered on GD4 delayed implantation without preventing pregnancy, though it resulted in smaller, less evenly spaced implantation sites and reduced implantation site weights (Kennedy, 1977). Furthermore, the selective PTGS2 inhibitors nimesulide and niflumic acid decrease ovulation rates in rats (Diao et al., 2007). While nimesulide given on GD4-8 had a modest effect on implantation, it altered key markers, including PPARδ, HB-EGF, and vimentin and impacted reproductive outcomes such as litter size, birth weight, and gestational length. (Diao et al., 2007). Celecoxib (PTGS2 inhibitor) and indomethacin (PTGS1 and PTGS2 inhibitor) significantly reduce pregnancy rates when given during GD 3-5 in rats (Sookvanichsilp and Pulbutr, 2002). Similarly, in mice, high doses of ibuprofen (PTGS1 and PTGS2 inhibitor) during the pre-implantation phase (GD0-3) resulted in implantation failure (Reese et al., 2001). Notably, concurrent PTGS1 and PTGS2 inhibition produces more severe reproductive effects than targeting either enzyme alone (Carp et al., 1988; Reese et al., 2001). Collectively, these findings across multiple species highlight the necessity of careful consideration when using NSAIDs during pregnancy (Supplementary Table 1).

As a non-selective PTGS inhibitor, indomethacin has been extensively used in reproductive research as a tool to investigate PG function during pregnancy (Kennedy, 1977; Kennedy, 1980; Lau et al., 1973; Saksena et al., 1976). Previous studies have established various dosing regimens and documented significant effects on reproductive outcomes. Early work by Lau et al. (1973) demonstrated that indomethacin at doses of 6-9 mg/kg (150-225 ug/day) could delay implantation in mice, while Kennedy (1977) showed that a much lower dose of 1mg/kg still resulted in delayed implantation in rats (Kennedy, 1977; Lau et al., 1973). However, these previous investigations primarily focused on binary pregnancy outcomes (implanted vs. non-implanted) and utilized conventional two-dimensional histological assessments. Thus although previous studies highlight the detrimental effects that NSAIDs have on pregnancy, the specific mechanisms by which NSAIDs alter the three-dimensional uterine architecture during peri-implantation stages, and how these structural changes translate to functional impairments are not well understood. We established a mouse model of pre-implantation indomethacin treatment followed by comprehensive three-dimensional analysis of implantation site architecture combined with temporal assessment throughout gestation. Our findings indicate that inhibiting both PTGS1 and PTGS2 together during the pre-implantation stage has a dose-dependent detrimental effect on pregnancy, affecting embryo spacing, on time implantation and implantation chamber formation, which in turn influences mid-gestation embryo resorption and litter size at birth.

## RESULTS

### Pre-implantation indomethacin treatment causes embryo crowding, delayed embryo implantation, compromised mid-gestation embryo development and reduced litter size at birth

First we established our NSAID model (Figure 1A) by intraperitoneal administration of 6mg/kg indomethacin (Lau et al., 1973) twice on gestational day (GD) 3 - at 0800h and 1400h. Embryos enter the uterine horn as a cluster around GD3 0000h and this administration regimen allows us to study the effect of NSAIDs on embryos that have already entered the uterine environment (Flores et al., 2020). Embryo implantation sites (EISs) at GD4.5 were visualized using intravenous blue dye injections. We observed that indomethacin-treated mice displayed 49% lower EISs than vehicle-treated mice (median blue dye sites number in vehicle: 14, 6mg/kg indomethacin: 8, P<0.0001) (Figure 1B, H). When we evaluated embryo number at GD3.75 by counting blastocysts in Hoechst-stained uteri (Flores et al., 2020) no differences were observed in embryo number between vehicle and 6mg/kg indomethacin treatment (Figure 1C, H). At the onset of decidualization at GD5.5 we observed embryo crowding in indomethacin treated mice, however the number of EISs were similar to vehicle treatment (Figure 1C) suggesting that the reduced number of EISs at GD4.5 was due to a delay in embryo implantation. We evaluated and observed a significant decrease in *Leukemia inhibitory factor* (*Lif*) expression in uterine glands at GD3.75, when compared to vehicle-treated uteri (Supplementary Figure 1A-C) that might explain the delay in implantation. 6 mg/kg indomethacin treatment did not affect uterine luminal epithelium closure at GD3.75 (Supplementary Figure 1D-E). Additionally, serum progesterone (P4) levels were comparable between vehicle and indomethacin treated mice (Supplementary Figure 1F), indicating that ovarian function was not impaired. Finally treatment of mice at GD3 with PTGS1-specific inhibitor (700nmol aspirin) or the PTGS2-specific inhibitor (600nmol Dup-697) did not delay embryo implantation or impact P4 levels (Supplementary Figure 2A-D).

**Figure 1.**
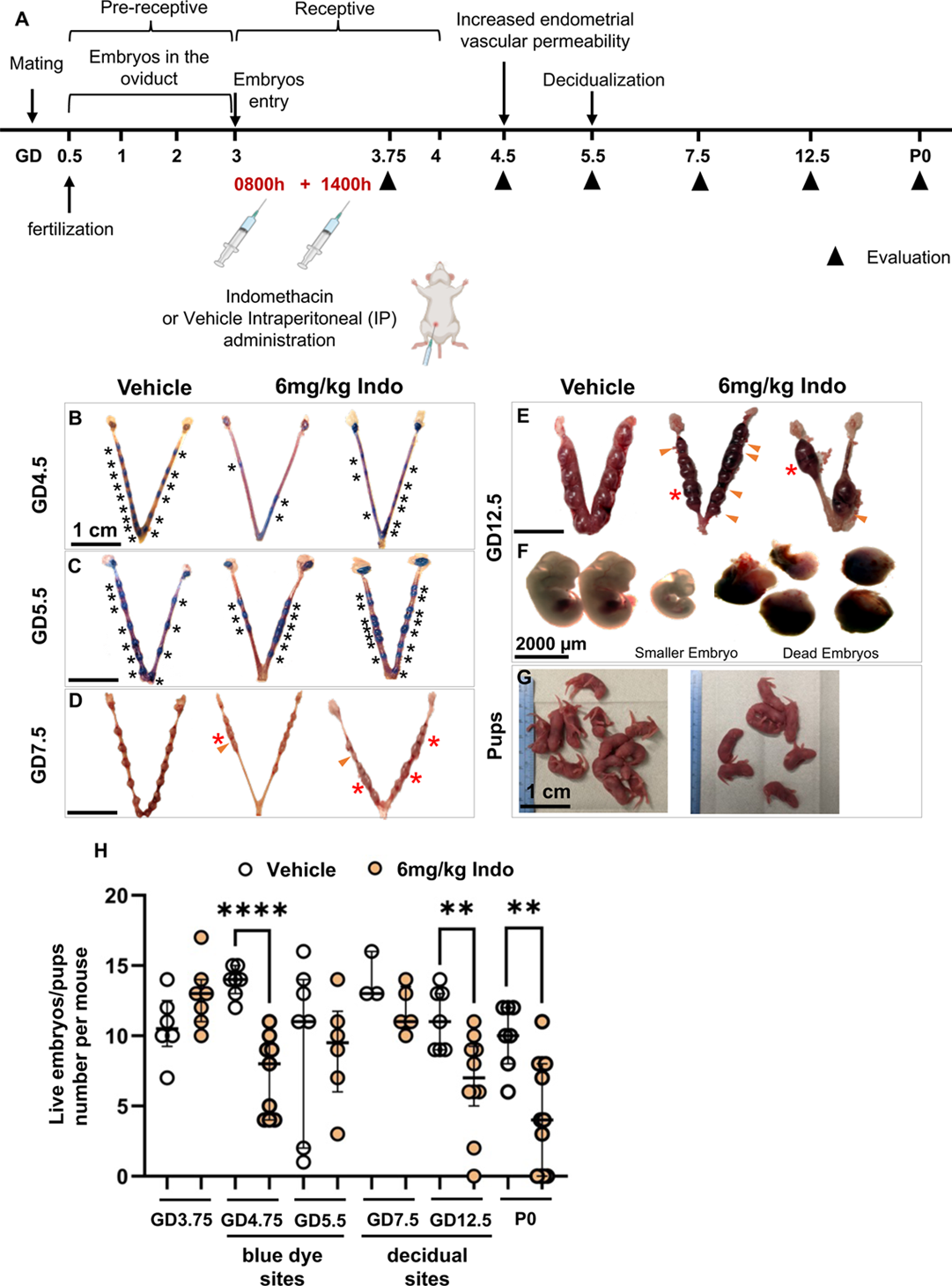
Pre-implantation indomethacin treatment delays embryo implantation, causes mid-gestation decidual resorption, and reduces litter size. **(A)** Timeline of mouse pregnancy, indicating indomethacin and vehicle treatment times and uteri evaluation throughout pregnancy. **(B, C)** Blue dye sites at GD4.5 and GD5.5. **(D, E)** Decidual sites at GD7.5 and GD12.5. **(F)** Embryos at GD12.5, illustrating small and dead embryos in 6mg/kg indomethacin-treated mice. **(G)** Litter born at P0 in vehicle and 6mg/kg indomethacin-treated uteri. Black asterisk: blue dye site; red asterisk: crowded decidua; orange arrowheads: resorbed decidua. Scale bars: B-E, G: 1 cm; F: 2000 µm. **(H)** Quantification of blue dye sites at GD4.5 and GD5.5, decidual sites at GD7.5 and GD12.5, and live pups at P0 in both treatment groups. At least n=3 mice per group. Each dot represents one mouse. Median values reported. Data analyzed using an unpaired parametric t-test. ** P < 0.01, **** P < 0.0001. Indo: indomethacin.

At GD7.5 we continued to observe embryo crowding and now also observed smaller decidual sites in uteri that were treated with indomethacin at GD3 (Figure 1D, H). For indomethacin treated mice, we observed 40% embryo resorption at GD12.5 (median live embryo number vehicle: 11, 6mg/kg Indo: 7, P<0.01) (Figure 1E, F, H), leading to a significant reduction in litter size at birth (median pups number in vehicle: 10, 6mg/kg Indo: 4, P<0.01) (Figure 1G, H). These effects of indomethacin treatment were dose dependent as treatment with 2mg/kg indomethacin at GD3 did not significantly affect the number of live pups born (Figure 1J, Supplementary Figure 3E). The low dose indomethacin did cause a significant reduction in blue dye sites at GD4.5 (median blue dye sites number in vehicle: 14, 2mg/kg indomethacin: 12, P<0.05) (Supplementary Figure 3B). However, in line with no impact on number of pups, the lower dose of indomethacin did not impact embryo number and embryo development at GD12.5 (Supplementary Figure 3B, C, D, E).

### Dose-dependent embryo crowding within indomethacin treatment

Since we observed embryo crowding, we further evaluated the effect of indomethacin on embryo distribution in the uterus. We determined the location of embryos along the longitudinal oviductal-cervical axis in the vehicle treated, 2 mg/kg indomethacin treated, and 6 mg/kg indomethacin treated mice (Figure 2A-C). We measured the distance each embryo traveled from the oviduct (oviduct-embryo (O-E) distance) and the embryo-embryo (E-E) distance at GD4.5 normalized for a uterine horn of 10 unit length. We observed no difference in the O-E distance across treatments (Figure 2A-D). However the E-E distance was significantly smaller in the 6mg/kg indomethacin treatment compared to both vehicle treatment and the 2mg/kg indomethacin treatment (the median E-E distance for vehicle: 1.17, 2 mg/kg Indo: 0.96, 6mg/kg indomethacin: 0.73, P < 0.01) (Figure 2A-D). We grouped embryos based on the normalized E-E distance and made 3 groups, E-E<0.5, 0.5<E-E<1 and E-E>1. We observed that in the 6 mg/kg indomethacin treatment group 33% of the embryos showed E-E<0.5, 33.81% were in the 0.5<E-E<1 group and 28.36% had E-E>1 (Figure 2D, E, Supplementary Table 2). In contrast, the majority of embryos in vehicle-treated uteri (75.64%) showed E-E>1 (Figure 2D, E, Supplementary Table 2). In the 2mg/kg Indomethacin group only 6% of the embryos showed E-E<0.5, 46% were in the 0.5<E-E<1 group and 48% had E-E>1 (Figure 2D, E, Supplementary Table 2). Thus, although the 2mg/kg showed a shift of E-E distances from >1 to 0.5<E-E<1, it appears that having E-E<0.5 in the 6mg/kg indomethacin treatment is what correlates with post-implantation embryo crowding.

**Figure 2.**
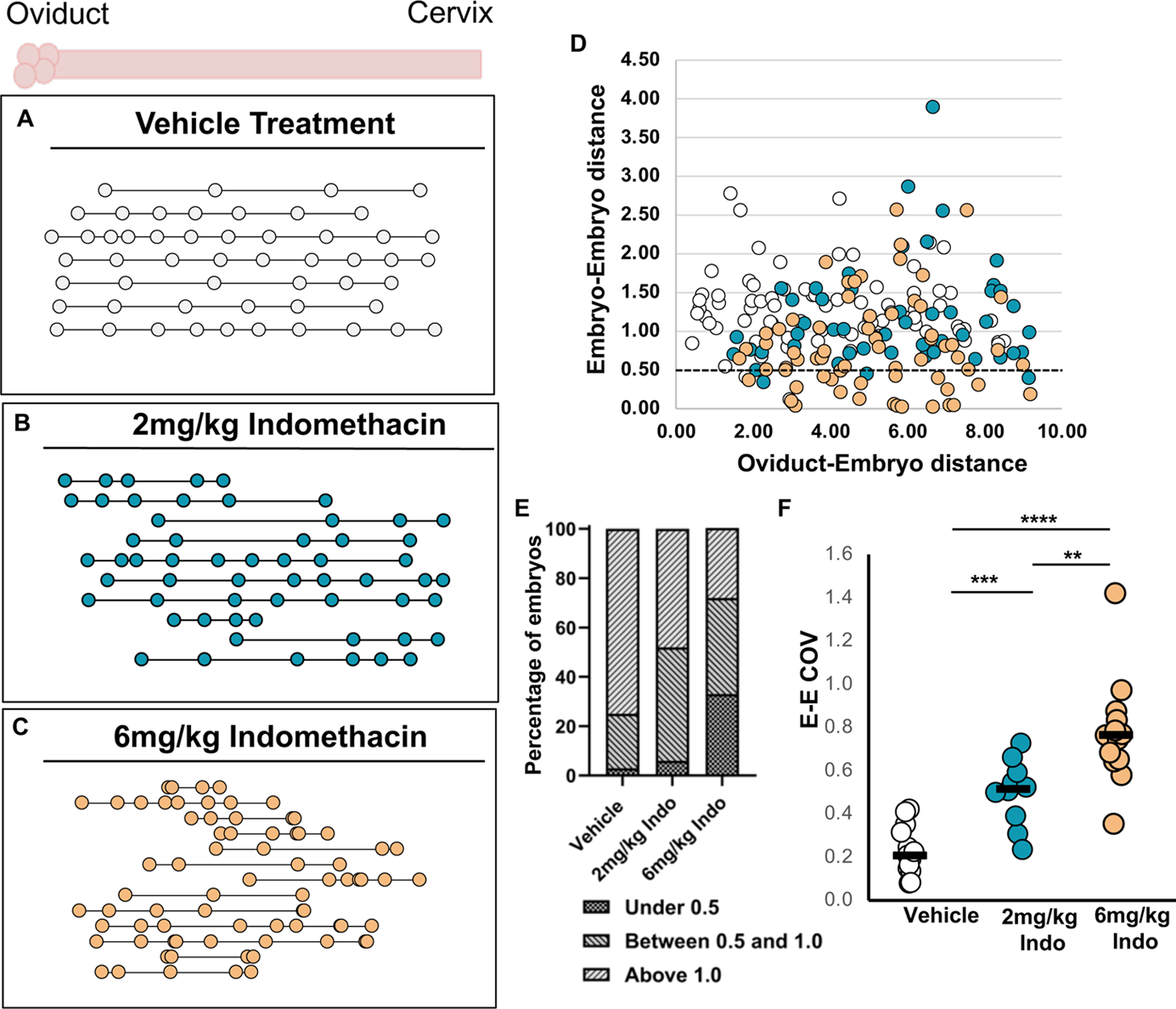
Uteri treated with 6mg/kg indomethacin show significant disturbance in embryo-embryo spacing compared to vehicle or 2mg/kg indomethacin treatments. (**A**) Placement of embryos along the oviduct-cervix axis in vehicle-treated mice, (**B**) 2mg/kg indomethacin-treated mice, and (**C**) 6mg/kg indomethacin-treated uterine horns at post-implantation stages on GD4.5. Each circle represents an individual embryo; circles connected by a line indicate embryos from the same uterine horn. The oviduct is on the left, cervix on the right. At least n=6 mice and 8 uterine horns analyzed per treatment group. (**D**) Classification of uterine horns based on oviduct-embryo (O-E) and embryo-embryo (E-E) distances. Each circle represents an embryo. Data analyzed using the Kruskal-Wallis test and Dunn’s multiple comparison test. ** P < 0.01, **** P < 0.0001. (**A-D**) White symbols: vehicle; light gray symbols: 2mg/kg indomethacin; black symbols: 6mg/kg indomethacin. (**E**) Percentage of embryos displaying the three ratios.: under 0.5 units, between 0.5-1.0 units, and above 1.0 units. Indo: indomethacin. (**F**) Coefficient of variation (COV) of E-E distances in vehicle, 2mg/kg and 6mg/kg indomethacin-treated uteri. Each dot represents one uterine horn. At least n=9 uterine horns per treatment were analyzed. Each dot represents one mouse. Median values shown. Data analyzed using an unpaired parametric t-test. **** P < 0.0001.

Next we calculated the coefficient of variation (COV) of E-E distances to determine how randomness of embryo distribution may be impacted. Vehicle-treated mice showed uniform embryo spacing with a mean COV of 0.227 ± 0.032, indicating non-random embryo spacing and proper embryo-uterine communication (Figure 2F) (Flores et al., 2020). Uteri treated with both 2mg/kg and 6mg/kg indomethacin exhibited a higher COV of 0.50 ± 0.048 and 0.775 ± 0.068 respectively suggesting an effect on embryo distribution in the second phase of movement (Figure 2F).

### Embryo development defects in indomethacin-treated uteri

To evaluate how pre-implantation inhibition of PTGS1 and PTGS2 might affect embryonic development we evaluated embryo quality at GD3.75 and GD4.5 and the epiblast stage at GD5.5. Embryos looked comparable in quality at GD3.75 and GD4.5 in vehicle and indomethacin treated uteri (Figure 3A-H, M). At GD5.5, ~48% of the embryos failed to develop into the epiblast stage. Some of these embryos were completely resorbed, others remained at the blastocyst stage, and the remainder did not attain the cylindrical shape characteristic of the epiblast (Figure 3I-L, M). To evaluate if embryo crowding was the sole contributor of observed developmental delays and defects in the embryos we separately analyzed crowded and uncrowded embryos. Our previous work has shown that peri-implantation regions (PIRs) at GD4.0 are >0.5 units in length for a horn of 10 unit length (Fig. 2C in (Madhavan et al., 2022)). Thus E-E distances <0.5 units would result in embryos attaching within the same PIR and could be a hallmark of embryo crowding. Given this finding, we separated the abnormal embryos at GD5.5 based on whether they were present in the same decidua (indicating crowding) or if they were present in separate decidua. ~42% of the abnormal-looking embryos were located in crowded sites (within the same decidua), while approximately 58% of the abnormally shaped embryos were present in distinct decidua (Figure 3N). Thus defective embryo development could be a result of crowding, delayed embryo implantation and a defective uterine environment.

**Figure 3.**
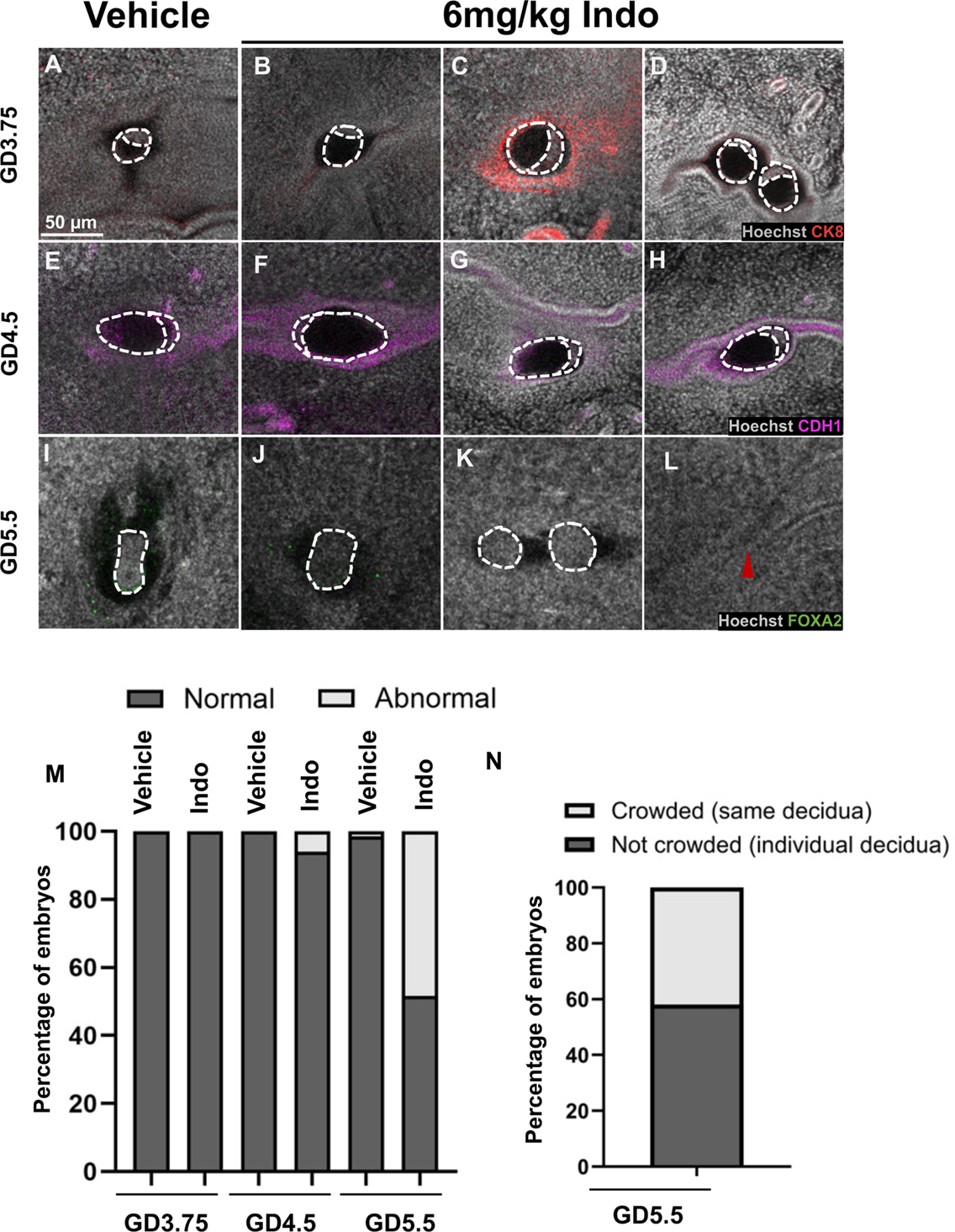
PTGS1 and PTGS2 inhibition restricts embryo growth at post-implantation stages on GD5.5. (**A**) Blastocyst-stage embryos in vehicle-treated mice at GD3.75. (**B-D**) Blastocyst-stage embryos in 6mg/kg indomethacin-treated mice at GD3.75. (**E**) Blastocyst-stage embryos in vehicle-treated mice at GD4.5. (**F-H**) Blastocyst-stage embryos in indomethacin-treated mice at GD4.5. (**I**) Epiblast-stage embryos in vehicle-treated mice at GD5.5. (**J-L**) Epiblast-stage embryos in 6mg/kg indomethacin-treated mice at GD5.5, showing abnormal epiblast shape, morula-stage embryos, and resorbed embryos. Dotted lines represent embryo shapes. Red arrowhead: resorbed embryo. At least n=6 mice per group analyzed per time point. Scale bars: A-H: 50 μm. Top of images: mesometrial pole; bottom: anti-mesometrial pole. (**M**) Comparison of normal vs abnormal embryo development from GD3.75 to GD5.5 in both treatment groups. (**N**) Percentage of abnormal embryos in crowded sites vs single decidua at GD5.5. Indo: indomethacin.

### Pre-implantation indomethacin treatment disrupts implantation chamber morphogenesis and vascular remodeling at post-implantation stages

Previously in a genetic deletion model we have shown that uterine deletion of PTGS2 results in resorption as early as GD4, 1800 hours (Massri and Arora, 2025) and these defects were due to poor implantation chamber growth. Thus we investigated implantation chamber formation in our indomethacin treated uteri. We observed that the length of the embryo implantation chamber was significantly reduced in the indomethacin-treated group at both GD4.5 (median implantation chamber length in the vehicle group: 515.0, in the 6 mg/kg indomethacin group: 447, P < 0.01) and GD5.5 (median implantation chamber length in the vehicle: 885µm, in the 6 mg/kg indomethacin group: 609µm, P < 0.0001) (Figure. 4A-I). We also found that blood vessel density surrounding the embryo implantation chamber was significantly elevated in indomethacin-treated uteri compared to their vehicle counterparts at GD4.5 (median blood vessel density in the vehicle: 21.03, versus in the 6 mg/kg indomethacin: 29.82, P < 0.05) (Fig. 4A, B, C). Analysis of CD31-positive cells revealed a clustering around the implantation chamber (Govindasamy et al., 2021), with this spatial distribution closely correlating to areas where PTGS2 is expressed (Massri and Arora, 2025). In our assessments, 100% of the implantation sites (22/22) in vehicle-treated mice exhibited a CD31 signal surrounding the implantation chamber (Fig. 4M-M’, Q), whereas only 29% (4/14) implantation sites in the mutant displayed a similar CD31 signal (Fig. 4N, N’, O, O’, Q). Thus pre-implantation indomethacin treatment results in shorter chambers with poor vessel development.

**Figure 4.**
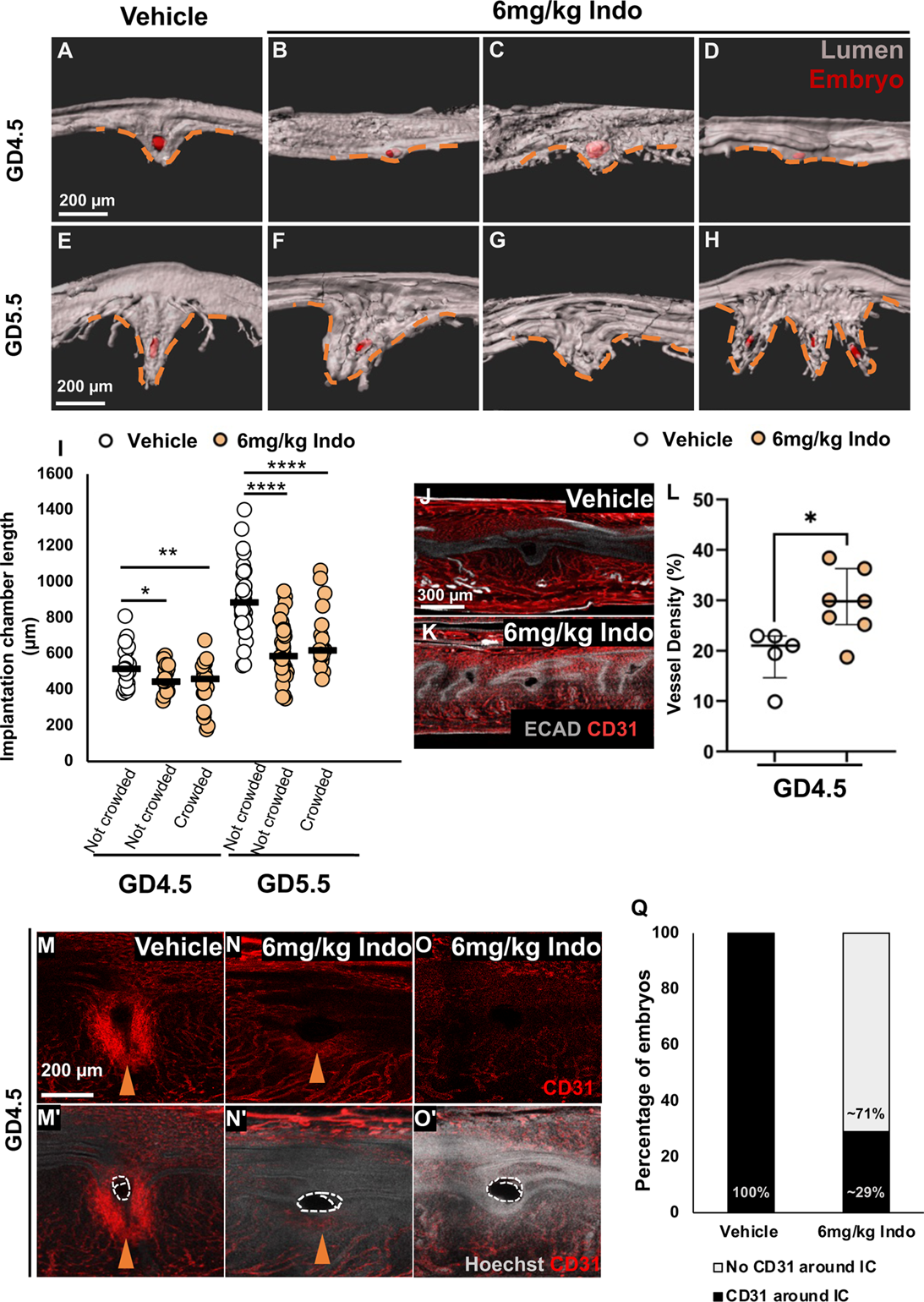
Abnormal embryo implantation chamber formation and length in uteri treated with 6 mg/kg indomethacin, and abnormal vascular development at embryo implantation sites in these uteri. (**A**) V-shaped implantation chambers in vehicle-treated mice at GD4.5. (**B-D**) Abnormally shaped implantation chambers in 6mg/kg indomethacin-treated mice at GD4.5. (**E**) Elongated implantation chambers in vehicle-treated mice at GD5.5. (**F-H**) Short and elongated implantation chambers in 6mg/kg indomethacin-treated mice at GD5.5. Orange dashed lines: embryo implantation chamber. Scale bars: A-H: 200 μm. Top of images: mesometrial pole; bottom: anti-mesometrial pole. (**I**) Quantification of implantation chamber length in vehicle and 6mg/kg indomethacin-treated uteri at GD4.5 and GD5.5. Embryos were separated for Indomethacin treatment as crowded or not crowded based on E-E<0.5 or >0.5 at GD4.5, and whether present within the same decidua or present in independent decidua at GD5.5. (**J**) Vessel architecture in vehicle-treated uteri at GD4.5. (**K**) Vessel architecture in 6mg/kg indomethacin-treated uteri at GD4.5. Scale bars: J-K: 300 μm. (**L**) Quantification of vessel density at embryo implantation sites in both groups at GD4.5. Median values shown. Each dot represents one implantation site. Data analyzed using an unpaired parametric t-test, * P < 0.05. (**M, M’**) CD31 expression around the embryo implantation chamber in vehicle-treated uteri. (**N-O’**) CD31 expression in 6mg/kg indomethacin-treated uteri. Orange arrowhead: CD31 expression, white dotted line: Embryo shape. Scale bars: M-O’: 200 μm. Top of images: mesometrial pole; bottom: anti-mesometrial pole. (**Q**) Quantification of embryo implantation chambers with and without CD31 expression. At least n= 4 mice per time point. Each dot represents one implantation chamber. Median values shown. Data analyzed using an unpaired parametric t-test and the Mann-Whitney test. *< P <0.05, *** P < 0.001, *** P < 0.0001.

## DISCUSSION

In agreement with prior literature (Kennedy, 1977; Kennedy, 1979; Kennedy et al., 2007; Lau et al., 1973) we show that pharmacological inhibition of PTGS1 and PTGS2 during the pre-implantation stage leads to embryo crowding, delayed embryo implantation, disruption of the embryo implantation chamber, and poor remodeling of blood vessels surrounding the implantation chamber. These disruptions appear to culminate in mid-gestation embryo resorption and a subsequent reduction in litter size.

### Delayed embryo implantation and disrupted embryo spacing are detrimental to pregnancy success

Delayed embryo implantation and disrupted embryo spacing can impair pregnancy success through distinct mechanisms (Diao et al., 2015; Song et al., 2002; Ye et al., 2005). Delayed embryo implantation may result in defective post-implantation development, manifesting as retarded embryonic growth and reduced litter size (Diao et al., 2007; Song et al., 2002; Song and Fazleabas, 2021). Transferring blastocyst-stage embryos to wild-type mice outside the implantation window causes significant anomalies and increased resorption rates (Song et al., 2002), emphasizing the critical importance of timing. Human studies also suggest that embryo implantation must occur within the window of implantation otherwise it could lead to an early pregnancy loss (Su and Fazleabas, 2015; Wilcox et al., 1999).

Embryo crowding can also compromise pregnancy success in multiparous species. During normal pregnancy, embryos space out evenly along the uterine horn to ensure adequate room for placental development and access to maternal resources (Chen et al., 2013; Flores et al., 2020). In genetic mutants where even spacing mechanism fails, as seen in Lpar3-deficient mice or Pla2g4a-deficient mice, embryos cluster together and implant at a single site (Song et al., 2002; Ye et al., 2005). This crowding event forces the embryos to compete for limited space and maternal resources, often resulting in shared placentas that cannot adequately support multiple embryos (Chen et al., 2013; Chen et al., 2011). This results in slower growth, nutrient restriction, and resorption, making proper spacing critical for pregnancy success (Chen et al., 2011; Ye et al., 2005). We have previously shown that embryos enter the uterus as a cluster and move to the center of the uterine horn by GD3 1200h. These embryos then scatter bidirectionally to space throughout the uterine horn (Flores et al., 2020). Our indomethacin treatment was performed after the embryos entered the uterus. This is likely why indomethacin treatment did not result in altered O-E distance. The clustering of embryos with reduced E-E distance in an indomethacin dose dependent manner observed in our study suggests that PTGS1/2-derived PGs are specifically essential for this bidirectional movement. PTGS2 signaling intersects with both LPAR3 and PLA2G4A signaling suggesting that these defects may be due to an effect of PGs on uterine contractility (Shah and Catt, 2005; Song et al., 2002; Ye et al., 2011).

Coefficient of Variation for the E-E distance is used to evaluate the distribution of embryos as a function of randomness (BOVING, 1956; Flores et al., 2020) or organized interaction between the embryos and the uterus. Previously we have shown that during the first phase of embryo movement when the embryos stay as a cluster and move unidirectionally, COV values are high suggesting the movement of the cluster is random. In the second phase of embryo spacing the COV values tend to decrease suggesting organized embryo-uterine interactions to allow for equal embryo spacing throughout the uterine horn. Higher E-E COV in indomethacin treated uteri confirm that embryos are stuck in their clustered movement stage and fail to switch over to the bidirectional scatter and spacing phase. This disruption has significant implications for the development of peri-implantation regions (PIRs). PIRs form through the resolution of uterine folds prior to the arrival of embryos at the implantation site, accompanied by gland reorientation toward the center of the which becomes the implantation site (Madhavan et al., 2022). During a normal pregnancy, the bidirectional movement of the embryos aligns with formation of PIRs, at least partially through a PG-dependent mechanism (Chen et al., 2013). The clustered embryos with E-E<0.5 suggests two possible failure modes: 1) multiple embryos clustering within a single PIR resulting in multiple implantation chambers within the same decidua as we observed. Even if these embryos elongate to the appropriate epiblast stage with suitable implantation chamber length, they will ultimately compete for decidual resources, which can lead to resorption. 2) Other embryos may become displaced into inter-implantation regions (IIRs) that lack fold resolution and embryos may die at the time of decidual expansion and shrinkage of the IIR regions. Both scenarios contribute to compromised peri-implantation success, explaining the increased rates of resorption and reduced litter sizes observed in our NSAID model.

### Indomethacin effects on post-implantation embryonic development

Our data indicate that pre-implantation inhibition of PTGS1 and PTGS2 disrupts the uterine environment essential for supporting embryonic development to the epiblast stage at GD5.5. The mechanisms underlying the observed defects in embryonic development likely involve both uterine and possibly embryonic factors. Our crowding analysis suggests that approximately 58% of abnormally developing embryos are situated in independent decidual sites, implying that the effects on development may be mediated both by the uterine environment and spatial competition among crowded embryos (Chida, 1985; van der Weiden et al., 1993). Furthermore, previous literature has highlighted that indomethacin can influence isolated embryonic development in vitro. Consequently, the systemic administration of indomethacin in our model may operate through several mechanisms. First, indomethacin disrupts uterine PG signaling, which is critical for creating a supportive environment for embryonic development, as evidenced by our findings regarding uterine-specific *Ptgs2* deletion (Massri and Arora, 2025). Second, indomethacin may induce mechanical crowding that limits individual embryos’ access to maternal resources. Finally, there is the possibility that indomethacin directly affects embryonic PG synthesis or signaling. Future studies employing wild type embryo transfer experiments—comparing indomethacin-treated and untreated females or transferring indomethacin-treated versus untreated embryos to control pseudo-pregnant mice—will help clarify the distinction between uterine-dependent and embryo-autonomous effects.

### Combined inhibition of PTGS1 and PTGS2 produces a more pronounced effect on pregnancy

Our results indicate that inhibiting PTGS1 and PTGS2 simultaneously produces more severe pregnancy defects than selective inhibition of either isoform alone (Supplementary Table 1). In our study, 700nmol Aspirin (PTGS1 specific inhibitor) and 600nmol Dup-697 (PTGS2 specific inhibitor) treatments at GD3, showed no signs of delayed implantation or embryo clustering at GD4.5 based on blue dye analysis. Although we did not extend these selective inhibitor experiments to assess term pregnancy events, the absence of peri-implantation defects contrasts sharply with the phenotypes caused by the non-selective PTGS inhibition caused by indomethacin. One factor that may contribute to the success of embryo implantation in cases of single isoform inhibition is a system of a backup mechanism that helps ensure reproductive success through supporting embryo implantation (Reese et al., 1999; Wang et al., 2004). The strongest evidence for this comes from genetic studies showing dramatically different outcomes when one versus both PTGS isoforms are eliminated. While mice lacking PTGS2 displayed complete implantation failure in C57BL/6J/129 mice (Lim et al., 1997), the same deletion in CD1 mice produces only partial reproductive defects because PTGS1 becomes upregulated in a pattern that mimics regular PTGS2 expression (Wang et al., 2004). The molecular basis for this compensatory upregulation in CD1 mice likely involves strain-specific polymorphisms in PTGS1 regulatory regions, enabling response to transcription factors such as NF-κB, C/EBP, or CREB, which typically regulate PTGS2 (Harper and Tyson-Capper, 2008; Kang et al., 2007). Alternatively, PG deficiency resulting from PTGS2 deletion could activate feedback loops that stimulate signaling pathways capable of inducing PTGS1 transcription in ways that resemble PTGS2 expression in endometrial tissues during peri-implantation stages (Chakraborty et al., 1996; Ricciotti and FitzGerald, 2011; Wang et al., 2004).

The critical importance of having at least one functional PTGS isoform is definitively demonstrated by dual PTGS1/PTGS2 knockout mice, which exhibit complete reproductive failure with no embryo attachment (Aikawa et al., 2024; Lim et al., 1997; Reese et al., 2001). Pharmacological studies support this conclusion: while selective inhibition of either PTGS1 or PTGS2 alone allows some reproductive function, simultaneous inhibition of both enzymes with non-selective NSAIDs like indomethacin or ibuprofen can prevent implantation (Supplementary Table 1, This study) (Kennedy, 1977; Lau et al., 1973; Reese et al., 2001). The clearest demonstration comes from Reese et al. (2001), who showed that even high-dose celecoxib (600 mg/kg twice daily, GD3-7) - a selective PTGS2 inhibitor - only modestly reduced PG levels and had minimal reproductive effects. However, combining this pharmacological PTGS2 inhibition with genetic PTGS1 deletion produced complete reproductive failure, demonstrating that simultaneous loss of both isoforms eliminates all PG sources in the female reproductive tract (Supplementary Table 1) (Reese et al., 2001). This phenotype is also in line with the complete implantation failure observed in PTGS1/PTGS2 double knockout mice (Aikawa et al., 2024). Another explanation for phenotype differences between some of the genetic models and pharmacological inhibition of PTGS1 and PTGS2 is extra-uterine sources of PTGS1 and PTGS2, such as endothelial cells and immune cells. Additionally, a systematic comparison between NSAIDs and genetic models would be beneficial to clarify critical distinctions in PTGS inhibition levels, duration of effects, and the precise timing of pregnancy failure. Future mechanistic studies will concentrate on the local versus systemic effects of PG signaling, investigate potential compensatory pathways in genetic models, and explore the non-PTGS effects of NSAIDs that may influence implantation outcomes.

### Limitations of the study

This study has some limitations. First, although we tested three different NSAIDs—indomethacin, DuP-697, and aspirin—that vary in their selectivity for PTGS1 and PTGS2, we did not measure PG levels in serum or uterine tissue during or after treatment. Without such measurements, we cannot definitively determine which PG (PGE2, PGF2α, PGI2, etc.) was affected by each drug or assess the actual inhibition levels of PTGS1 versus PTGS2 in vivo at the doses used. This limits our ability to establish clear mechanistic links between specific prostaglandin pathways and the adverse pregnancy outcomes observed. Second, in our study we considered crowding as embryos within the same decidual site. It is unclear whether embryos that are in independent decidua but in closer proximity to other decidua would compromise pregnancy outcomes. Third, using only one animal model (mice or rats) may not fully recapitulate human pregnancy physiology, particularly regarding PTGS expression patterns and prostaglandin-dependent processes during implantation.

## Conclusion

Our study suggests that NSAID administration during the critical window of embryo implantation disrupts embryo spacing and the three-dimensional structure of the post-implantation chamber, which contributes to the reduction in litter size and leads to unfavorable pregnancy outcomes. Therefore, caution should be exercised when administering NSAIDs during early pregnancy, considering our results and other studies that indicate a significant disruption to the maternal environment that ultimately compromises its ability to support embryonic growth and development.

## MATERIALS AND METHODS

### Animals

In this study, we utilized adult CD1 mice aged 6 to 10 weeks. For the pregnancy experiments, adult females of the same age were paired with fertile CD1 males. The presence of a vaginal plug indicated gestational day 0 (GD0) at 1200h. We euthanized mice at various stages, specifically at GD3.75 (on GD3 between 1800h and 2100h), GD4.5, GD5.5, GD7.5, and GD12.5, (on respective GD between 1200h to 1500h). Additionally, some mice were allowed to reach term to assess litter size. For GD4.5 and GD5.5 dissections, animals were euthanized 10 minutes following a 0.15 ml intravenous injection of 1.5% Evans blue dye (MP Biomedicals, ICN15110805). The uteri from GD4.5 to GD12.5 were photographed under white light to document implantation and decidual sites. All mice were kept on a 12-hour light/dark cycle, and all studies and protocols received approval from the Institutional Animal Care and Use Committee at Michigan State University.

### Drugs

Indomethacin (Sigma-Aldrich, I7378-5G) was dissolved in 0.01% Dimethyl sulfoxide (DMSO) in PBS to a final concentration of 1mg/ml, and mice were intraperitoneally injected with either 150 ul or 50 ul (6mg/kg or 2mg/kg) on gestational day (GD) 3 at 0800h and 1400h. Aspirin (Sigma-Aldrich, # PHR1003) was dissolved in 0.01% ethanol in PBS to a final concentration of 7 mM, and 100 μl (700 nmol) (Lim et al., 1997) of this solution was intraperitoneally injected in CD1 mice on GD3 at 0800h and 1400h. Dup697 (Enzo Life Sciences, #BMLEI3580005) (PTGS2 specific inhibitor) was dissolved in 0.1% DMSO in PBS to a final concentration of 6 mM. Mice were intraperitoneally injected with 100 μl (600 nmol) (Lim et al., 1997) intraperitoneally on GD3 at 0800h and 1400h. The drug doses were selected based on previous studies (Lau et al., 1973; Lim et al., 1997). Control mice for all drug treatments received only vehicle treatment (0.01% DMSO in PBS, 0.01% ethanol in PBS, or 0.1% DMSO in PBS, respectively) on GD3 at 0800h and 1400h. The twice-daily dosing schedule was chosen based on the pharmacokinetic properties of these compounds (Bindu et al., 2020).

To assess the clinical relevance of these dosages, human equivalent doses (HEDs) were calculated using the FDA-recommended body surface area scaling method: HED = mouse dose × (mouse Km/human Km) = mouse dose × (3/37). For aspirin, the 700 nmol dose (approximately 126 μg per 20g mouse, or ~6.3 mg/kg) corresponds to an HED of 0.51 mg/kg, equivalent to approximately 36 mg for a 70 kg adult, which is within the low-dose aspirin range used clinically (75-100 mg daily for cardioprotection). For indomethacin, the 6 mg/kg dose corresponds to an HED of 0.49 mg/kg (approximately 34 mg for a 70 kg adult), which falls within the standard therapeutic range of 25-50 mg taken two to three times daily, with maximum daily doses of 150-200 mg. Dup697, being a selective research compound without established human therapeutic doses, was used at concentrations previously validated in mouse studies for PTGS2 inhibition (Lim et al., 1997). This allometric scaling method effectively considers the differences in metabolic rate and body surface area between species, making it more physiologically relevant than merely scaling by body weight. The calculated HED indicates that the mouse dosage used is clinically pertinent and would be expected to produce therapeutic effects similar to those experienced by human patients receiving standard indomethacin treatment. A limitation is that our doses were administered intraperitoneally and the human dosage is for oral consumption.

### Whole-mount immunofluorescence staining

For whole-mount staining of dissected uteri, we initiated the process by fixing the samples in a cold mixture of DMSO and methanol in a 1:4 ratio as previously described (Arora et al., 2016; Flores et al., 2020; Madhavan et al., 2022; Massri and Arora, 2025). To ensure adequate hydration, the specimens were immersed in a 1:1 solution of methanol and PBST (PBS with 1% Triton) for 15 minutes, followed by a thorough wash in 100% PBST for another 15 minutes. Subsequently, the samples were placed in a blocking solution composed of PBS, 1% Triton, and 2% powdered milk for 1 hour at room temperature. After this, we proceeded with the incubation of the samples in primary antibodies (Supplementary Table 3) mixed in the blocking solution for seven nights at 4°C. Following the primary antibody incubation, the samples underwent a washing phase, which involved two washes of 15 minutes and then four washes of 45 minutes each with 100% PBST. We then introduced Alexa Fluor-conjugated secondary antibodies, incubating the samples for three nights at 4°C as specified in Supplementary Table 3. Afterward, the samples were washed again with 100% PBST—first with two washes of 15 minutes and then four washes of 45 minutes. To further prepare the samples, we incubated them at 4°C overnight in a solution of 3% H2O2 diluted in methanol. The final steps involved washing the samples with 100% methanol for a specified duration and then clearing the tissues overnight with a benzyl alcohol and benzyl benzoate solution in a 1:2 ratio (Sigma-Aldrich, 108006, B6630).

### Cryo-embedding, cryo-sectioning, and immunostaining

As described previously (Massri and Arora, 2025) we fixed uterine tissues in 4% paraformaldehyde (PFA) for 20 minutes and subsequently incubated the samples in fresh 4% PFA overnight at 4°C. The tissues were washed with 100% PBS three times for five minutes each, followed by overnight incubation in a 10% sucrose/PBS solution at 4°C. We then transferred the samples to 20% and 30% sucrose solutions in PBS, allowing them to sit for 2-3 hours each at 4°C. After that, we embedded the samples in Tissue-Tek OCT (Andwin Scientific, 45831) and stored them at −80°C. Cryo-sections measuring 7µm in thickness were mounted onto glass slides (Fisher, 1255015). For immunofluorescent staining, we let the slides air dry for 15 minutes before washing them with 100% PBS three times for five minutes each, then blocked them with a solution of PBS, 2% powdered milk, and 1% Triton for 20 minutes. After performing another round of washes with 100% PBS three times for five minutes, we stained the slides with primary antibodies (Supplementary Table 3) and incubated them at 4°C overnight. The following day, we washed the slides with 100% PBS three times for five minutes and treated the sections with secondary antibodies and Hoechst (Supplementary Table 3) for one hour at room temperature. Finally, after washing with PBS, we applied two drops of 20% glycerol in PBS to the slides and then sealed the sections with glass coverslips.

### In situ hybridization

We conducted in situ hybridization on uterine sections with the RNAscope 2.5 HD Assay-RED kit (ACD Bio, 322350), which also features immunofluorescence capabilities, as previously outlined (Granger et al., 2024; Massri and Arora, 2025). Our goal was to identify *Lif* mRNA linked to the uterine glands at GD3.75. To detect *Lif*, we employed the Mm-Lif probe (ACD Bio, 475841), and for labeling the uterine glands, we performed immunostaining for FOXA2 (Supplementary Table 3). The complete 3-day protocol was executed following the guidelines established by ACD Bio (322360-USM, MK 51-149 TN).

### Serum progesterone measurement

After euthanizing the mouse, we collected blood samples ranging from 200 to 500 µl, allowing them to sit at room temperature for 30 minutes. Next, we centrifuged the samples for 15 minutes at 2000 g, carefully separated the supernatant, and stored the samples immediately at −20°C. Once the samples were collected and preserved, we sent them to the Ligand Assay and Analysis Core Laboratory in Charlottesville, VA, for progesterone level analysis. The samples were diluted in a 1:4 ratio, tested in triplicate for accuracy, and the results were reported in ng/ml.

### Confocal microscopy

We used a Leica SP8 TCS white light laser confocal microscope utilizing 10x air to image whole uterine tissues or 20X water objective and a 7.0 um Z stack or system-optimized Z stack to image the samples (Madhavan et al., 2022). Upon imaging, we imported the files (.LIF format) into Imaris v9.2.1 (Bitplane; Oxford Instruments, Abingdon, UK) 3D surpass mode. We created 3D renderings using surface modules.

### Image analysis

#### Implantation chamber, luminal epithelium, and embryo visualization

We used the CDH1 fluorescent signal for the luminal epithelium surface and the FOXA2 fluorescent signal for uterine glands to visualize the implantation chamber. We isolated the luminal epithelium by subtracting the FOXA2-specific signal from the CDH1 signal. We used the Hoechst signal to locate embryos based on the inner cell mass (ICM) signal, and we used the 3D rendering surface in IMARIS software to create the embryo surfaces. We used the measurement function in Imaris to measure the length of the implantation chamber from the mesometrial to the anti-mesometrial pole.

#### Leukemia inhibitory factor (Lif) quantitation

As previously detailed (Granger et al., 2024; Massri and Arora, 2025), we utilized the FOXA2 signal to create 3D surfaces of the nuclei in the glands using the 3D surface function within IMARIS software. We then employed the IMARIS masking function to generate a separate channel for the Lif signal, which is situated beneath the 3D surface of the uterine glands that was established earlier. Utilizing this new channel for the *Lif* signal, we developed an additional 3D surface for *Lif*. After constructing the 3D surfaces, we measured the 3D surface volume of both the glands and *Lif* using the statistics function in Imaris. Finally, we calculated the *Lif* volume in relation to the uterine gland volume using Microsoft Excel and plotted this data as *Lif* volume per uterine gland volume (FOXA2 signal) with normalized units. volume (FOXA2 signal) with normalized units.

#### Vessel density around the anti-mesometrial pole of the implantation chamber

We developed a 3D rendering surface that visualizes blood vessels by employing a CD31 fluorescent signal. In addition, we created a channel in Imaris software to mask the blood vessel surfaces around the embryo implantation sites. For effective image segmentation, we imported 14 µm of the masked vessel channel into ImageJ after adjusting the scale and utilizing the threshold function. By applying the vessel analysis Plugin in ImageJ (https://imagej.net/), we assessed the density of blood vessels at the embryo implantation sites. Vessel density data is represented as the percentage of area occupied by blood vessels as shown previously (Massri and Arora, 2025).

#### Embryo location

To perform a comprehensive analysis of embryo locations, we initiated the process by creating 3D renderings of key anatomical features: the oviductal-uterine junction, the embryos themselves, and the uterine horns as described previously (Flores et al., 2020). This step was accomplished using the Surface module, which allowed us to visualize these structures in three dimensions effectively. Following the creation of these renderings, we utilized the Measurements module to identify and record the three-dimensional Cartesian coordinates for the center of each surface. By projecting these coordinates onto the x-y plane, we were able to calculate: the distance between the oviductal-uterine junction and each embryo (referred to as O-E), the distances between adjacent embryos (designated as E-E), and the overall length of the uterine horn. To address the natural variations in uterine horn lengths observed among the different mice, we normalized all measured distances to the respective horn length. This approach allows us to present the mean values for O-E and E-E as ratios, ensuring they are unitless and comparable across samples.

### Statistical analysis

We used Microsoft Excel and GraphPad Prism (Dotmatics; GraphPad, La Jolla, CA, USA) to analyze the statistical differences between the treatment groups and plot our graphs. To analyze the difference between the two treatment groups, we employed the unpaired parametric two-tailed t-test. First, we tested the data for homogeneity of the variance between the two treatments. If the variances were equal, we proceeded with a parametric two-tailed t-test. If the variances differed, we used the Mann-Whitney U-test to compare the two treatment groups. Due to heterogeneity of variances between the three groups, differences between treatment groups (vehicle control, low dose, and high dose indomethacin) were analyzed using the Kruskal-Wallis non-parametric test. When the Kruskal-Wallis test indicated significant differences (p < 0.05), post-hoc comparisons were performed using Dunn’s multiple comparison test to identify specific group differences. We considered the data statistically different for P value < 0.05 or less.

## ACKNOWLEDGEMENTS

We thank Dr. Asgerally Fazleabas, Dr. Nataki Douglas, Dr. Shuo Xiao and Dr. Gregory Burns for critical discussions related to the project.

## AUTHOR CONTRIBUTIONS

NM and RA conceptualized the study and designed the experiments. NM and SW performed the experiments. NM, ML and RA analyzed and validated the data. NM and RA prepared the figures and wrote and edited the manuscript. All authors reviewed and accepted the final version of the manuscript.

---

**Supplementary Figure 1.**
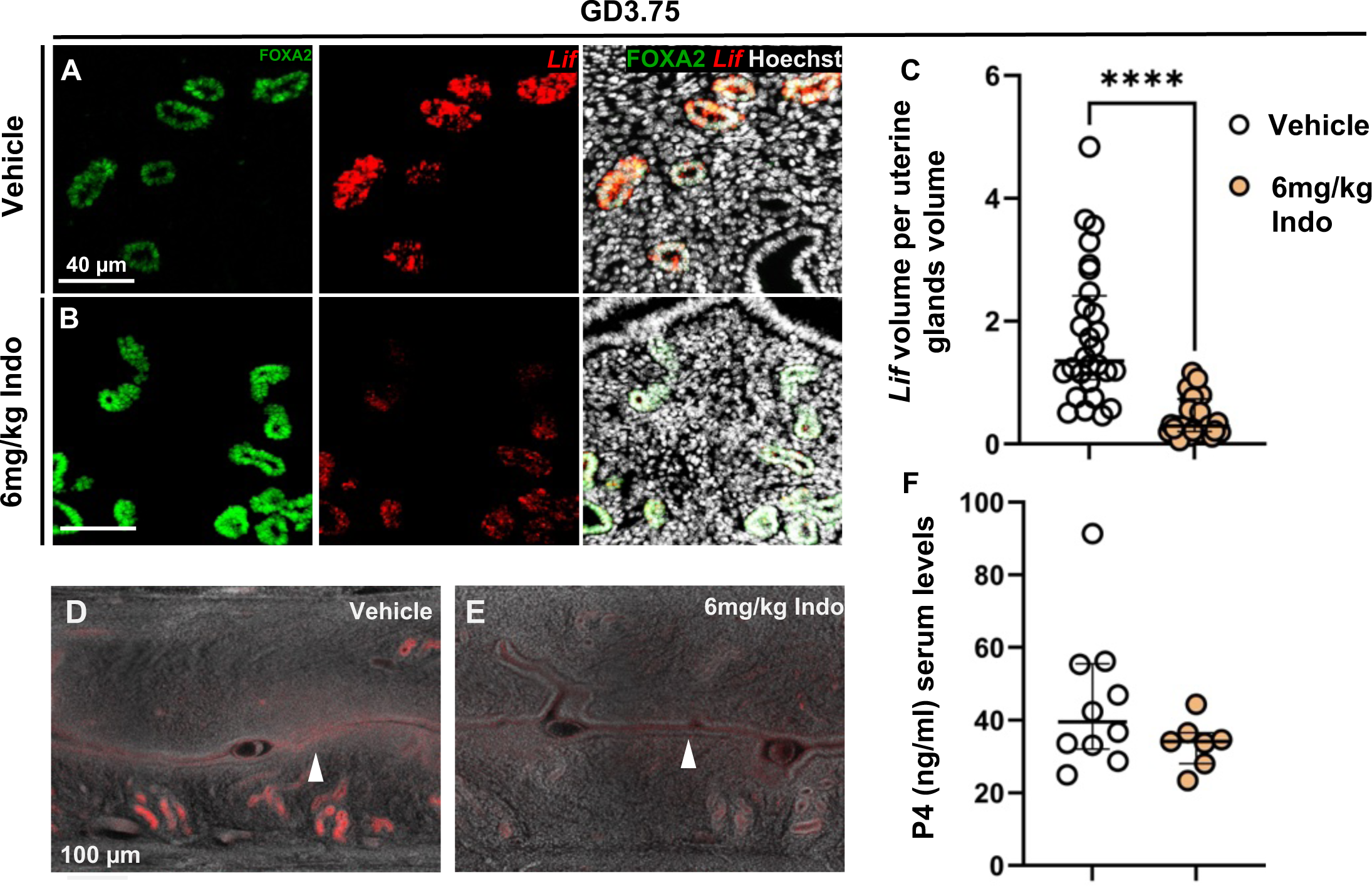
6mg/kg indomethacin-treated uteri display reduced preimplantation Leukemia Inhibitory Factor (Lif) expression with no effect on luminal closure or progesterone levels. **(A, B)** Lif expression in FOXA2+ glandular epithelium cells in vehicle and 6mg/kg indomethacin-treated uteri at GD3.75. **(C)** Quantification of Lif volume normalized to FOXA2+ glandular epithelium volume at GD3.75 per uterine section in both groups. At least n=4 mice, and 28 7µm sections analyzed per group. Each dot represents one uterine section. Median values shown. Data analyzed using the Mann-Whitney test. **** P < 0.0001. **(D)** Luminal closure representation in vehicle-treated mice. **(E)** Luminal closure in 6mg/kg indomethacin-treated mice at GD3.75. **(F)** Progesterone serum levels in both groups at GD3.75. At least n=3 mice per treatment analyzed. Each dot represents one mouse. Median values shown. Data analyzed using an unpaired parametric t-test. No significant difference observed.

**Supplementary Figure 2.**
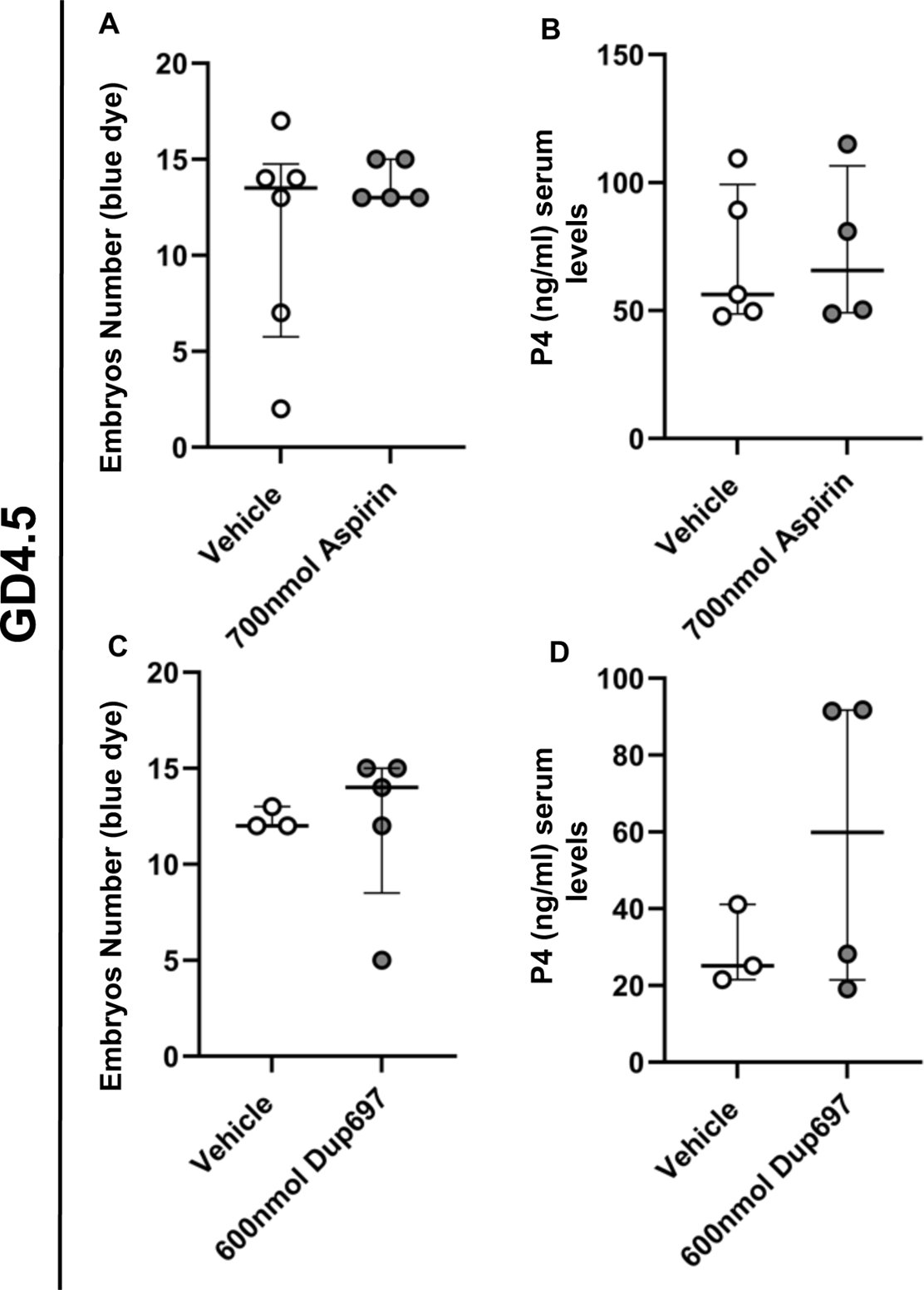
Pre-implantation treatment of 700nmol aspirin or 600nmol dup-697 does not affect implantation success. **(A)** Quantification of embryo number at GD4.5 in vehicle and 700nmol aspirin-treated groups. At least n=3 mice per group. Each dot represents one mouse. Median values reported. Data analyzed using an unpaired parametric t-test. **(B)** Progesterone serum levels in both groups at GD4.5. At least n=3 mice per treatment were analyzed. Each dot represents one mouse. Median values shown. Data analyzed using an unpaired parametric t-test. No significant difference observed. **(C)** Quantification of embryo number at GD4.5 in vehicle and 600nmol Dup-697-treated groups. At least n=3 mice per group. Each dot represents one mouse. Median values reported. Data analyzed using an unpaired parametric t-test. **(D)** Progesterone serum levels in both groups at GD4.5. At least n=3 mice per treatment were analyzed. Each dot represents one mouse. Median values shown. Data analyzed using an unpaired parametric t-test. No significant difference observed.

**Supplementary Figure 3.**
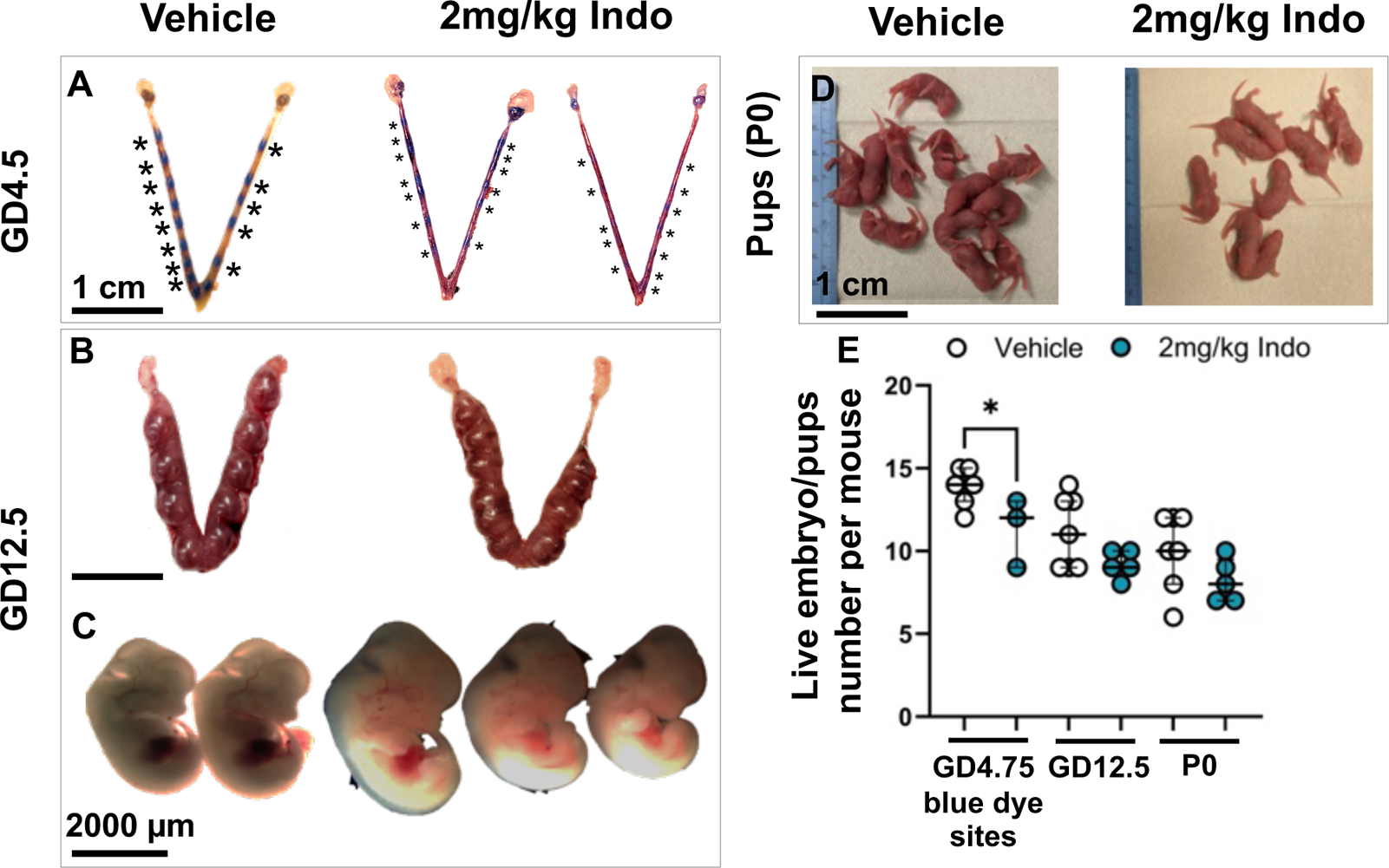
Pre-implantation treatment of 2mg/kg indomethacin does not affect overall pregnancy success. **(A)** Blue dye sites at GD4.5. **(B)** Decidual sites at GD12.5. **(C)** Embryos at GD12.5. **(D)** Litter size at P0 in vehicle and 2mg/kg indomethacin-treated uteri. Black asterisk: blue dye site. Scale bars: A, B, D: 1 cm; C: 2000 µm. **(E)** Quantification of embryo number at GD4.5, GD12.5, and live pups at birth (P0) in vehicle and 2mg/kg indomethacin-treated groups. At least n=3 mice per group. Each dot represents one mouse. Median values reported. Data analyzed using an unpaired parametric t-test. * P < 0.05. Indo: indomethacin.

**Table 1:**
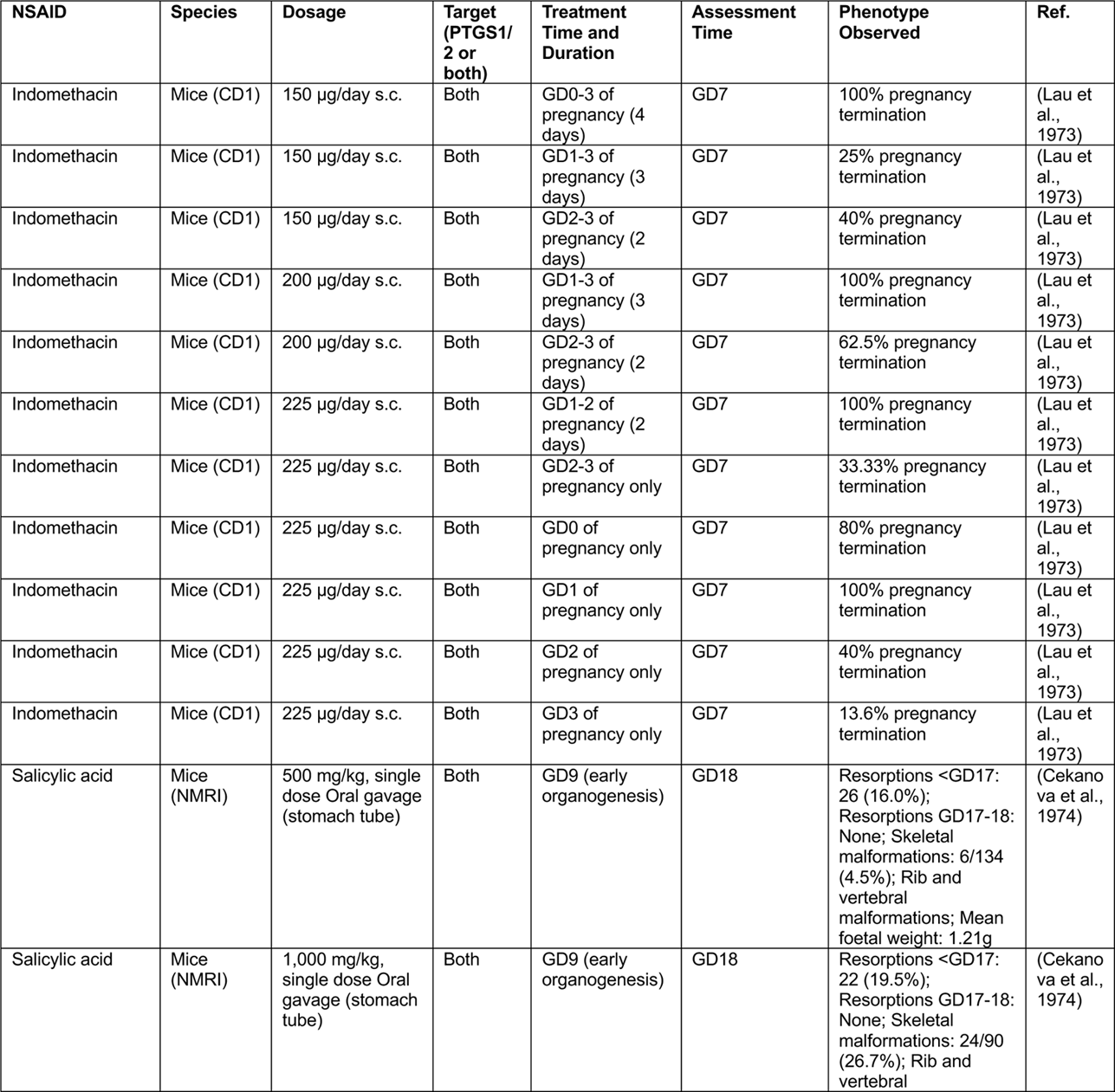
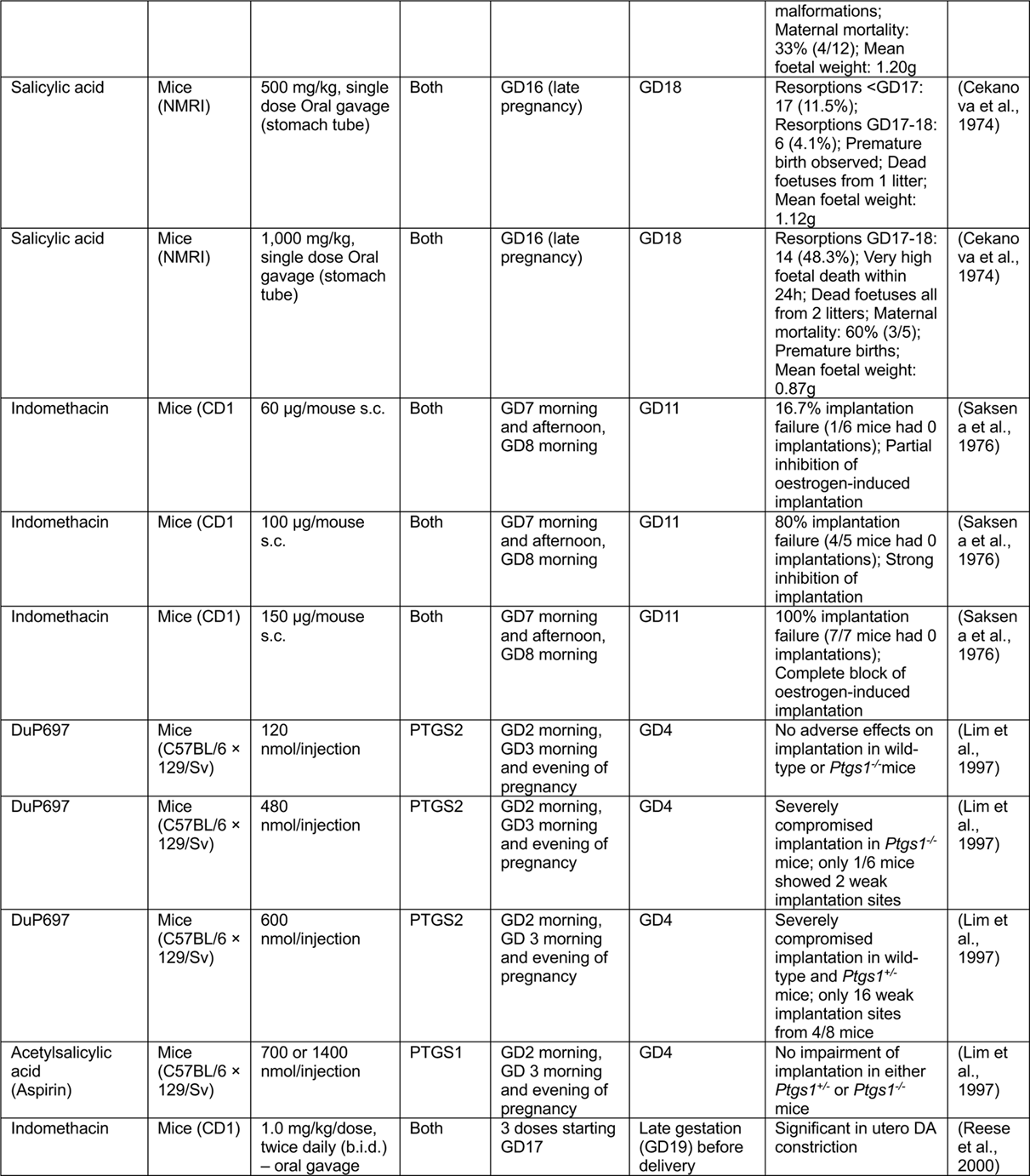
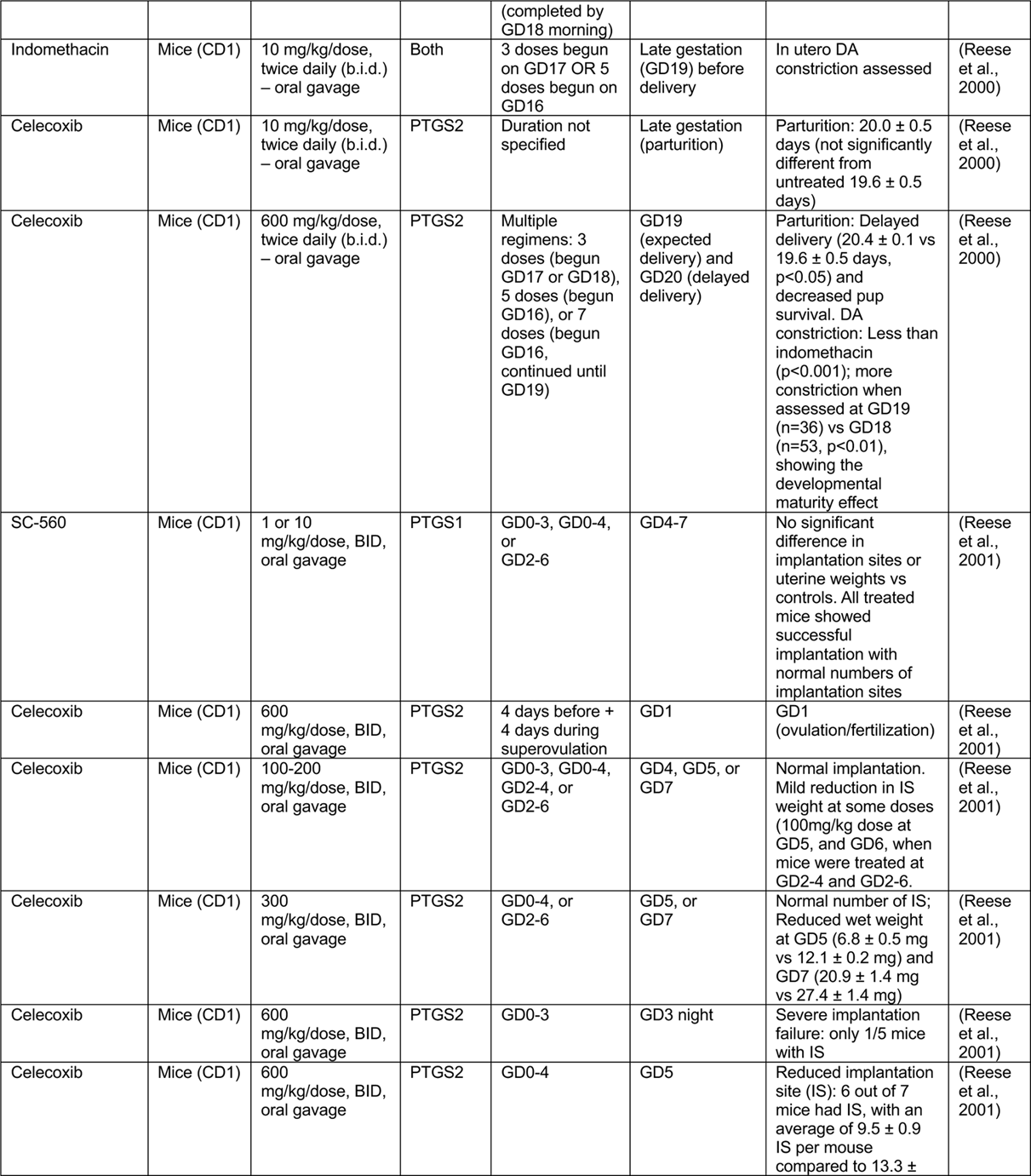
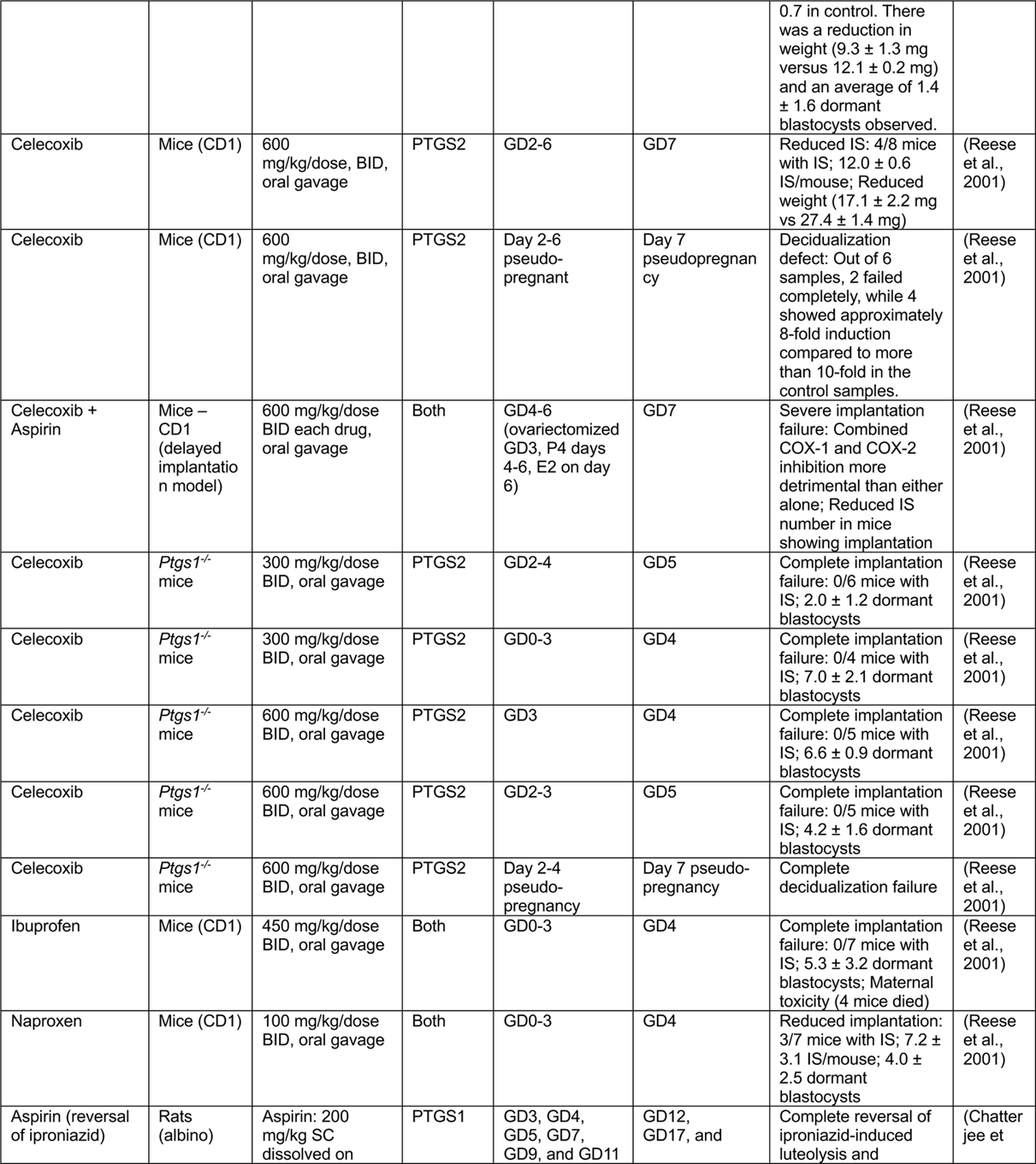
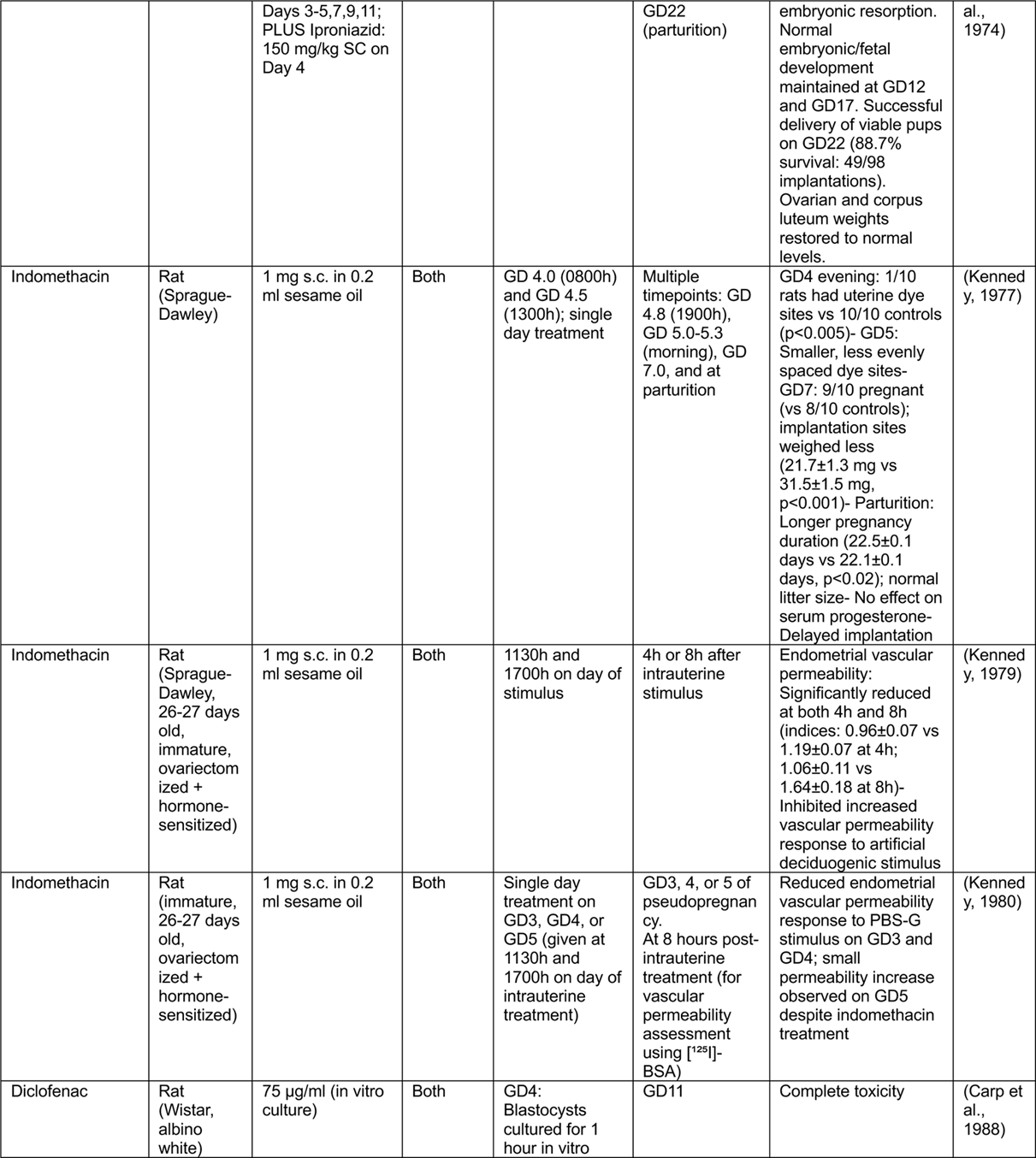
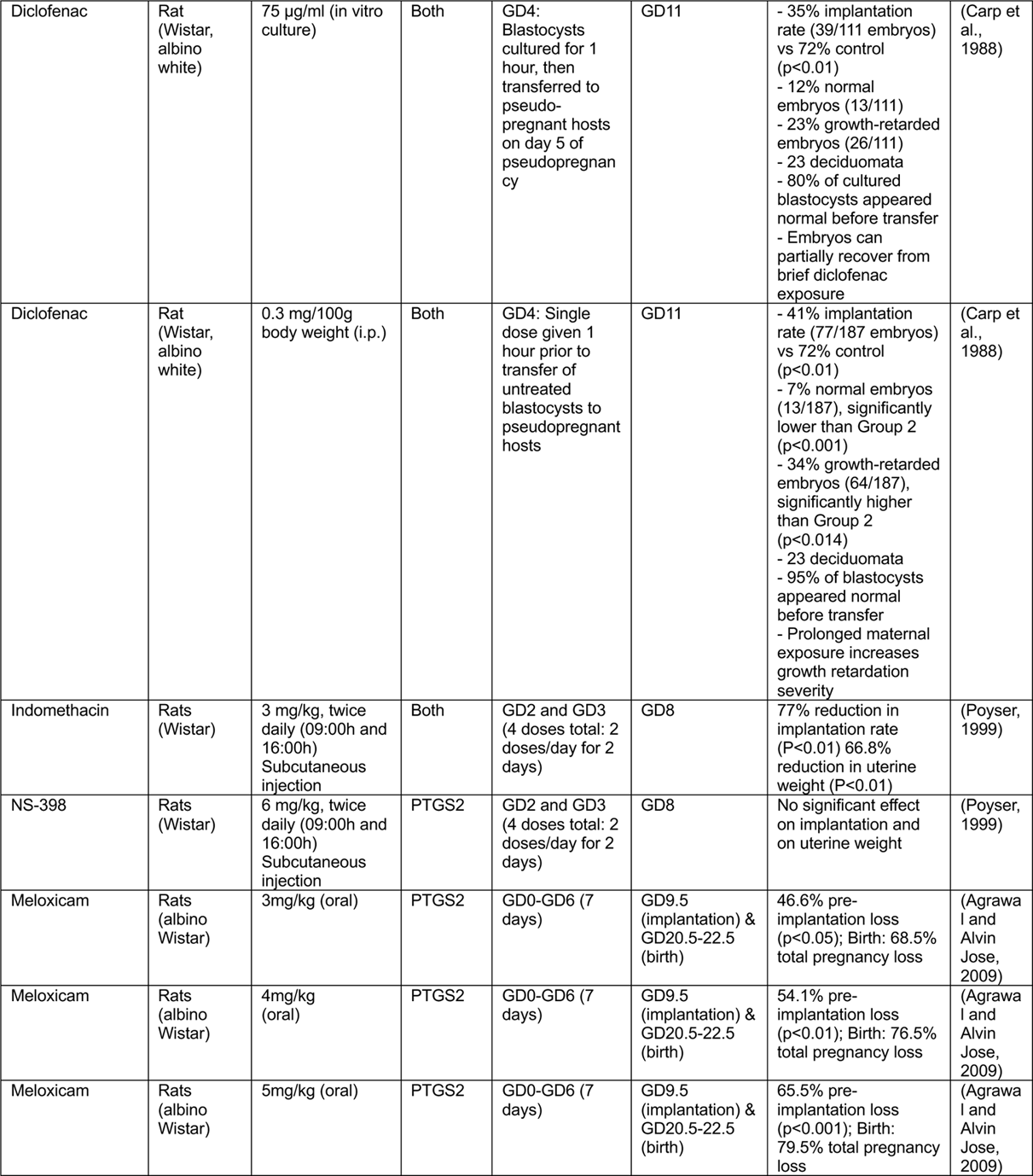
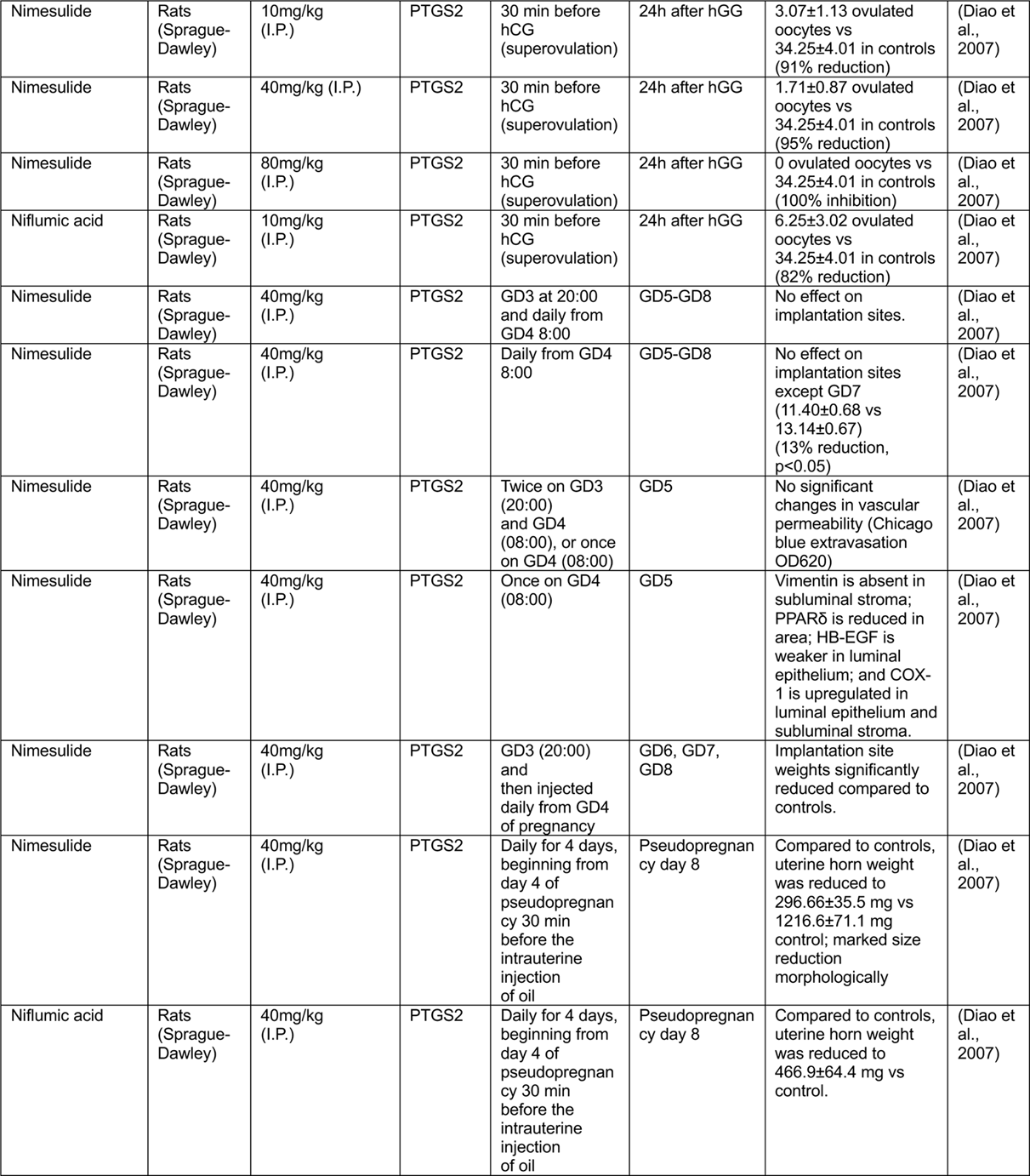
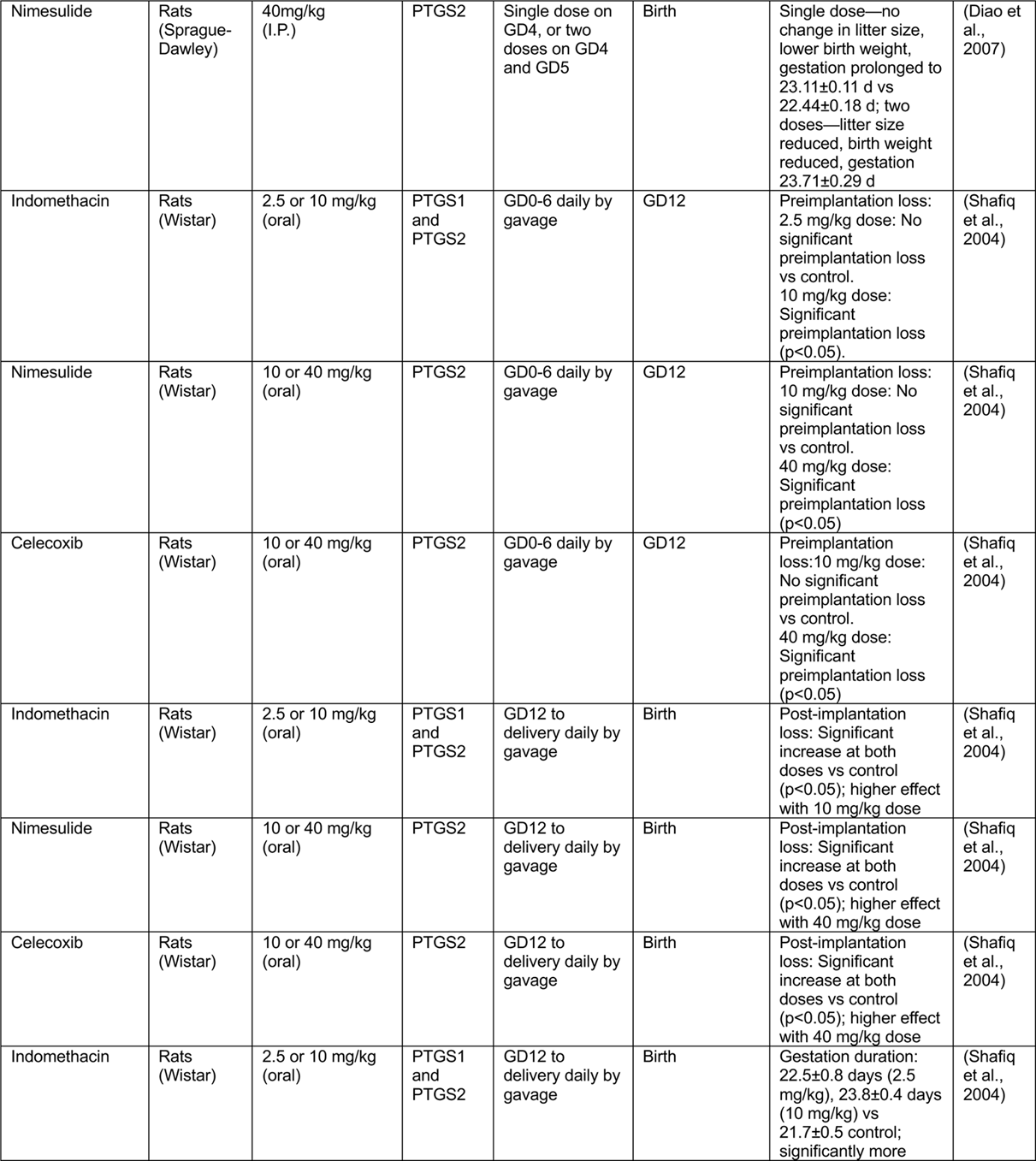
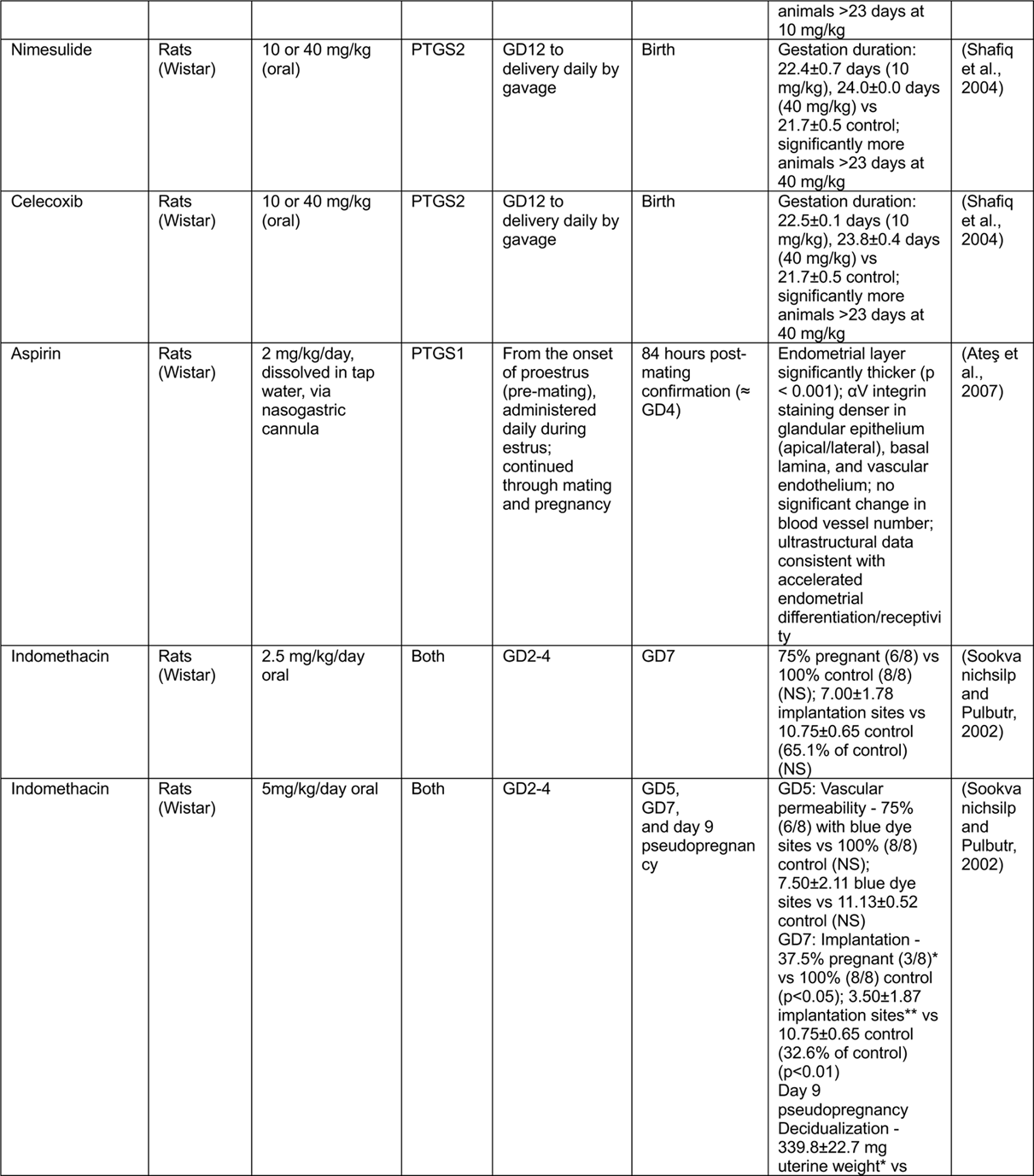
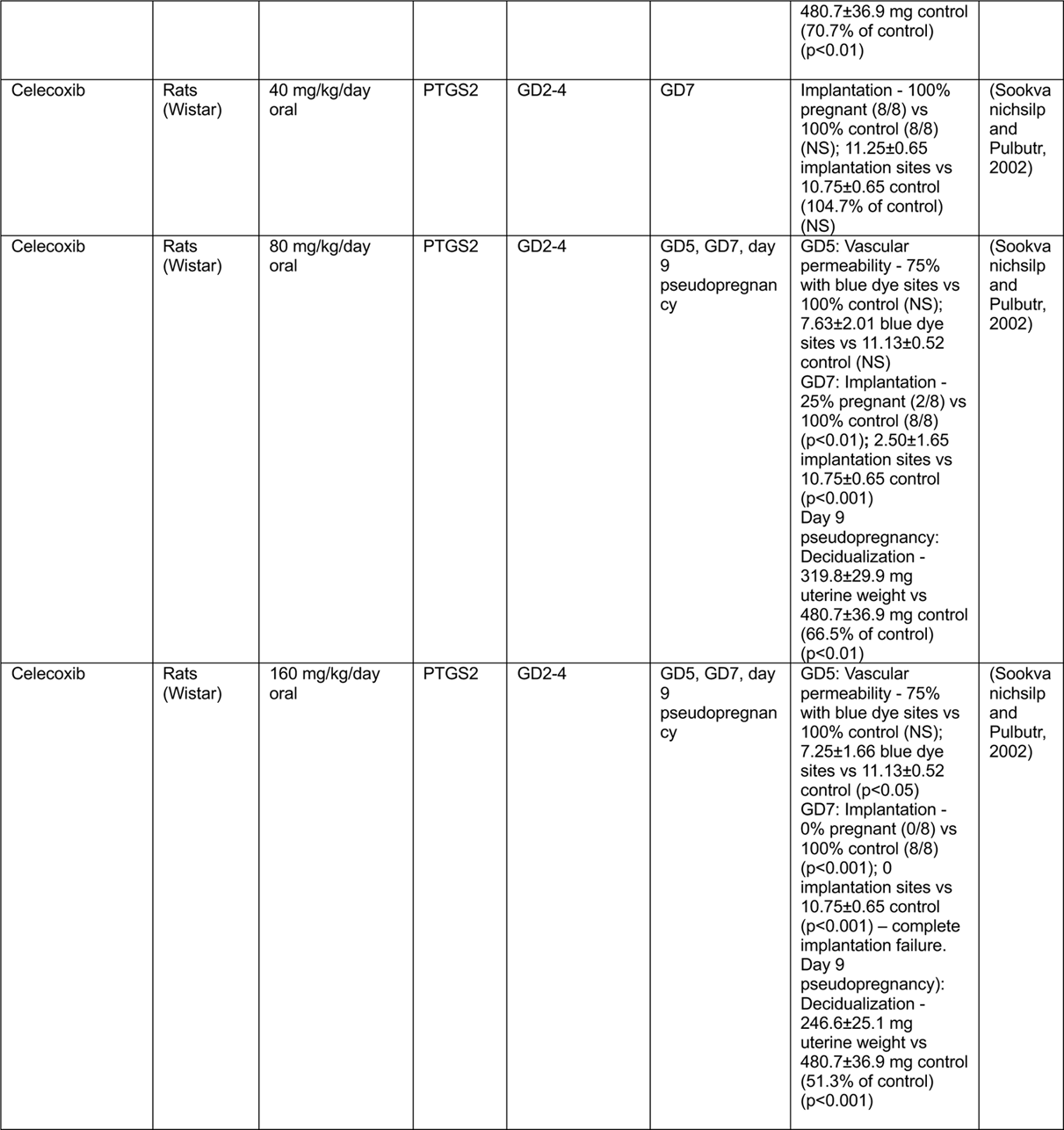
Summary table for NSAIDs used in rodents (mice/rats) with the following columns showing the name of NSAIDs, species, dosage, and administration methods, target, treatment time and duration, assessment time, and phenotype observed. GD: gestational day. Day of plug: GD0.

**Table 2:** Distribution of embryo-embryo distance along the oviduct-cervical axis considering the horn length to be 10 units. This table presents the distribution of embryo-embryo distance in different ratios (< 0.5, 0.5-1.0, > 1.0) across three treatment groups: Vehicle, 2mg/kg Indomethacin, and 6mg/kg Indomethacin. The data shows the number of embryos (n) and the percentage of the total embryos within each range group.

|  | **< 0.5** | **0.5-1.0** | **> 1.0** |
| --- | --- | --- | --- |
|  | **n (%)** | **n (%)** | **n (%)** |
| **Vehicle** | 2 (2.56) | 17 (21.79) | 59 (75.64) |
| **2mg/kg Indo** | 3 (6) | 23 (46) | 24 (48) |
| **6mg/kg Indo** | 22 (32.84) | 26 (38.81) | 19 (28.36) |

**Table 3:** Primary and secondary antibodies used in the study.

| **Primary antibody** | **Dilution** | **Stained tissues** |
| --- | --- | --- |
| Rat-anti-CDH1 (M108, 108006, B6630) | 1:500 | Luminal epithelium |
| Rabbit anti-FOXA2 (Abcam, ab108422) | 1:200 (sections) | Glandular epithelium |
| Armenian-hamster anti-CD31 (DSHB, AB_2161039) | 1:200 | Endothelial cells |
| **Secondary antibody** | **Dilution** |  |
| Goat anti-Armenian hamster 647 (Invitrogen, A78967) | 1:500 |  |
| Goat anti-rat 647 (Invitrogen, A21247) | 1:500 |  |
| Donkey anti-rabbit 555 (Invitrogen, A31572) | 1:500 |  |
| **Nuclear marker** | **Dilution** |  |
| Hoechst (Sigma Aldrich, B2261) | 1:500 |  |

